# Overcoming resistance to immune checkpoint therapy in PTEN-null prostate cancer by sequential intermittent anti-PI3Kα/β/δ and anti-PD-1 treatment

**DOI:** 10.1101/2020.10.17.343608

**Authors:** Zhi Qi, Zihan Xu, Liuzhen Zhang, Yongkang Zou, Wenyu Yan, Cheng Li, Ningshu Liu, Hong Wu

**Affiliations:** The MOE Key Laboratory of Cell Proliferation and Differentiation, Peking University, Beijing 100871, China; School of Life Sciences, Peking University, Beijing 100871, China; Peking-Tsinghua Center for Life Sciences, Peking University, Beijing 100871, China; Bayer AG, Drug Discovery TRG Oncology, Muellerstrasse 178, 13353 Berlin, Germany

**Keywords:** Overcoming resistance to ICT, PTEN-null prostate cancer, intermittent PI3K inhibitor treatment, anti-PD-1 therapy, cancer cell-intrinsic immunosuppressive activity, tumor microenvironment, tertiary lymphoid structures, CD8^+^ clonal expansion and memory, CD8^+^ T cell-mediated anti-tumor immunity

## Abstract

Prostate cancers generally lack T cell infiltration and display resistance to immune checkpoint therapies (ICT). We found that intermittent but not daily dosing of PI3Kα/β/δ inhibitor BAY1082439 on a *Pten*-null spontaneous prostate cancer model could overcome ICT resistance and unleash CD8^+^ T cell-dependent anti-tumor immunity *in vivo*. Mechanistically, BAY1082439 converts *Pten*-null cancer cell-intrinsic immune-suppression to immune-stimulation by promoting IFNα/γ pathway activation, β2-microglubin expression and CXCL10/CCL5 secretion. Together with its preferential Treg inhibition activity, BAY1082439 promotes clonal expansion of tumor-associated CD8^+^ T cells. Once primed, tumors remain as T cell-inflamed and become responsive to anti-PD-1 therapy. Our data suggest that intermittent PI3K inhibition can alleviate *Pten*-null cancer cell-intrinsic immunosuppressive activity and turn “cold” tumors into T cell-inflamed ones, paving the way for successful ICT.

**Significance:** The combination of ICT and targeted therapies holds great promises for broad and long-lasting therapeutic effects for cancers. However, combining ICT with anti-PI3K inhibitors have been difficult because the multifaceted effects of PI3K on both cancer cells and immune cells within the tumor microenvironment. Here we show a carefully designed anti-PI3K treatment, both in its specificity and dosing schedule, to inhibit cancer cell growth while promoting anti-tumor immunity, is critically important for successful ICT. Since the PI3K pathway is one of the most frequently altered signaling pathways in human cancers, our work may shed light on treating those cancers with PI3K activation and overcome resistance to ICT.

**Highlights:**

- Intermittent PI3Kα/β/δ inhibitor BAY1082439 treatment overcomes ICT resistance
- BAY1082439 turns *Pten*-null prostate cancer from “cold” to T cell-inflamed
- BAY1082439 inhibits cancer cell-intrinsic immunosuppressive activity and Treg
- BAY1082439 promotes clonal expansion and immunity of tumor-associated CD8^+^ T cells

## Introduction

Immune checkpoint therapies (ICT), such as those mediated by anti-PD-1 or CTLA-4 antibodies, have shown promising long-lasting effects on certain cancer types by activating T cell-mediated anti-tumor immunity (Sharma and Allison, 2015; Sharma and Allison, 2020). The efficacy of ICT is positively correlated with the density of tumor infiltrating CD8^+^ T cells (Daud et al., 2016; Peng et al., 2016; Spranger et al., 2015; Tumeh et al., 2014). However, the majority of solid cancers have poor CD8^+^ T cell infiltration (“cold” tumors) and do not respond to ICT (Jansen et al., 2018; Zou et al., 2016). Although the mechanisms underlying cancer-mediated T cell exclusion are largely unknown, it has become clear that promoting T cell infiltration may increase the range of cancers sensitive to ICT (Bonaventura et al., 2019) (Galluzzi et al., 2018; Joyce and Fearon, 2015).

Prostate cancer is the most common malignancy in males, and the second leading cause of male cancer-related death in the Western world (Siegel et al., 2020). Androgen deprivation therapy (ADT) is the mainstream treatment for prostate cancer. However, despite initial regression, many patients progress to highly aggressive castration-resistant prostate cancer (CRPC), a disease stage with limited treatment options (Watson et al., 2015). Recent ICT trials on CRPC patients have shown disappointing results (Jansen et al., 2018; Sharma et al., 2020; Taghizadeh et al., 2019; Topalian et al., 2012), most likely due to low mutational load and defects in T cell-mediated anti-tumor immunity. It has been reported that over 90% of prostate cancers are “cold” and do not express a T cell-inflamed gene signature (Cristescu et al., 2018). Therefore, treatments that can promote T cell infiltration may pave the way for efficient ICT on prostate cancers.

Loss of the *PTEN* tumor suppressor or activation of its controlled PI3K pathway are associated with resistance to ICT in multiple tumor types (George et al., 2017; Peng et al., 2016). In prostate cancer, *PTEN* mutations have been found in 40-50% primary and 70-90% metastatic tumors (Taylor et al., 2010). PTEN loss in the murine prostatic epithelial in the *PB-Cre^+^Pten^L/L^* (*Pten*-null) mouse model can mimic both molecular and pathological features associated with human prostate cancers, including upregulated PI3K pathway, invasive adenocarcinoma, as well as resistant to ADT (Jiao et al., 2007; Mulholland et al., 2011; Wang et al., 2003). Cancer cell-intrinsic PI3K activation in the *Pten*-null model also promotes an immune suppressive microenvironment, including increased immune suppressive myeloid-derived suppressor cells (MDSC) and regulatory T cells (Treg), decreased dendritic cell maturation, as well as decreased T cell infiltration and activation (Calcinotto et al., 2018; Garcia et al., 2014; Tang et al., 2012). These results suggest that inhibiting cancer cell-intrinsic immune suppressive activity induced by PI3K activation may be a prerequisite for promoting T cell infiltration and achieving anti-tumor immunity.

However, targeting PI3K pathway is not a simple task, especially to achieve synergy with ICT, as PI3K pathway plays significant roles on both sides of the aisle. As one of the most important oncogenic pathways, PI3K activation promotes cell proliferation, survival, migration, angiogenesis, metabolic reprograming as well as an immune suppressive environment (Cheng et al., 2020; Okkenhaug et al., 2016; Sivaram et al., 2019; Thorpe et al., 2015; Wu et al., 2019); on the other hand, PI3K is also a critical regulator for the functions of immune cells within the tumor microenvironment, and inhibition of PI3K activity in these immune cells may be detrimental for ICT (Carnevalli et al., 2018; De Henau et al., 2016; Kaneda et al., 2016; Lu et al., 2017). To make things even more complicated, there are four isoforms in the class I PI3K family: PI3Kα/β are ubiquitously expressed and often abnormally activated in cancer cells by constitutively activated *PI3KCA* mutations or loss-of-function *PTEN* mutations, while PI3Kδ/γ are commonly restricted to leukocytes and essential for immune surveillance. Various immune cells within the tumor microenvironment are preferentially relying on different isoforms of the PI3K to promote or inhibit tumor development (Bilanges et al., 2019; Jia et al., 2008; Thorpe et al., 2015; Wang et al., 2015).

We recently reported that, BAY1082439, a new, selective PI3K inhibitor with equal potency against the PI3Kα/β/δ isoforms, was highly effective in inhibiting primary and CRPC in the *Pten*-null model (Zou et al., 2018). However, the effects of BAY1082439 on alleviating cancer cell-intrinsic immunosuppressive activity and on various types of immune cells within the tumor microenvironment, particularly CD8^+^ T cells, have not been investigated. In this study, we report that by changing the dosing schedule from daily to intermittent, BAY1082439 can generate favorable anti-tumor immune response through alleviating cancer cell-intrinsic immunosuppressive activity, directly inhibiting Treg cells, promoting IFNα/γ pathway activation and CD8^+^ cell infiltration and clonal expansion. As a consequence, intermittent treatment of BAY1082439 paved the way for effective ICT therapy.

## Results

### *Pten*-null prostate cancers are poorly T cell infiltrated and resistant to anti-PD-1 immunotherapy

Most prostate cancers are immunogenically “cold” with low number of infiltrating T cells although ADT is known to promote T cell infiltration (Cristescu et al., 2018; Jansen et al., 2018). We confirmed this clinical observation on the *Pten*-null prostate cancer model. Immunohistochemistry (IHC) staining of prostate tissues from the *Pten*-null mice showed that CD8^+^ T cells were scarcely present in the intact mice but increased upon castration (Fig. S1A). However, most of the T cells remained in the stroma area and could not penetrate into the tumor acini even after castration (Fig. S1A).

We then sought to test if T cells-induced upon castration could lead to antitumor immunity in the presence or absence of ICT. Castrated *Pten*-null mice were treated with either the anti-PD-1 or isotype control antibody for 4 weeks (Fig. S1B). FACS analysis revealed that although tumor-associated CD8^+^ T cells and Treg cells were slightly increased upon anti-PD-1 treatment, the CD8^+^ /Treg ratio remained unchanged in comparison to the control antibody (Fig. S1C). Hematoxylin and eosin (HE) and IHC assessments further confirmed that the numbers and localization of CD8^+^ and Granzyme b^+^ (GZMb^+^) cells after anti-PD-1 therapy remained unchanged (Fig. S1D). Therefore, the *Pten*-null prostate cancer model mimics human prostate cancers with poor T cell infiltration and resistance to anti-PD-1 monotherapy. *Pten*-null prostate cancer model could serve as an *in vivo* model to investigate the molecular mechanism underlying the T cell exclusion phenotype-associated with human prostate cancers.

### BAY1082439 inhibits the cancer cell-intrinsic growth and immunosuppression and activates IFNα and IFNγ pathway

We have shown in our previous study that PTEN-loss in the prostatic epithelial cells promotes immunosuppressive microenvironment during prostate cancer initiation and progression but the underlying mechanisms are largely unknown (Garcia et al., 2014). We hypothesized that PTEN-loss or PI3K pathway activation in cancer cells may modulate anti-cancer immune response, which dampens the immune activities of cells within the tumor microenvironment.

We treated *Pten*-null prostate cancer-derived cell lines CAP2 and CAP8 (Jiao et al., 2007) and *PTEN*-null human prostate cancer cell line PC3 and LNCaP with BAY1082439 to investigate the effects of BAY1082439 on cancer cell-intrinsic properties. As expected for PI3K pathway inhibition, global gene expression analysis demonstrated that BAY1082439 treatment led to downregulation of 9 mTOR signaling and cell proliferation-related pathways in all prostate cancer cell lines (Fig. 1A; Supplementary Table 2). Interestingly, the interferon α and γ (IFNα and IFNγ) response pathways were the 2 commonly upregulated pathways found in BAY1082439 treated CAP2/CAP8/PC3 lines but not in LNCaP line, as LNCaP lacks JAK1 expression, which is required for IFNα and IFNγ pathway activations (Dunn. et al., 2005) (Fig. S2A). Importantly, treating the *Pten*-null prostate cancer *in vivo* model with a bullet dose of BAY1082439 showed similar effects (Fig. 1A; Supplementary Table 2).

**Figure 1.**
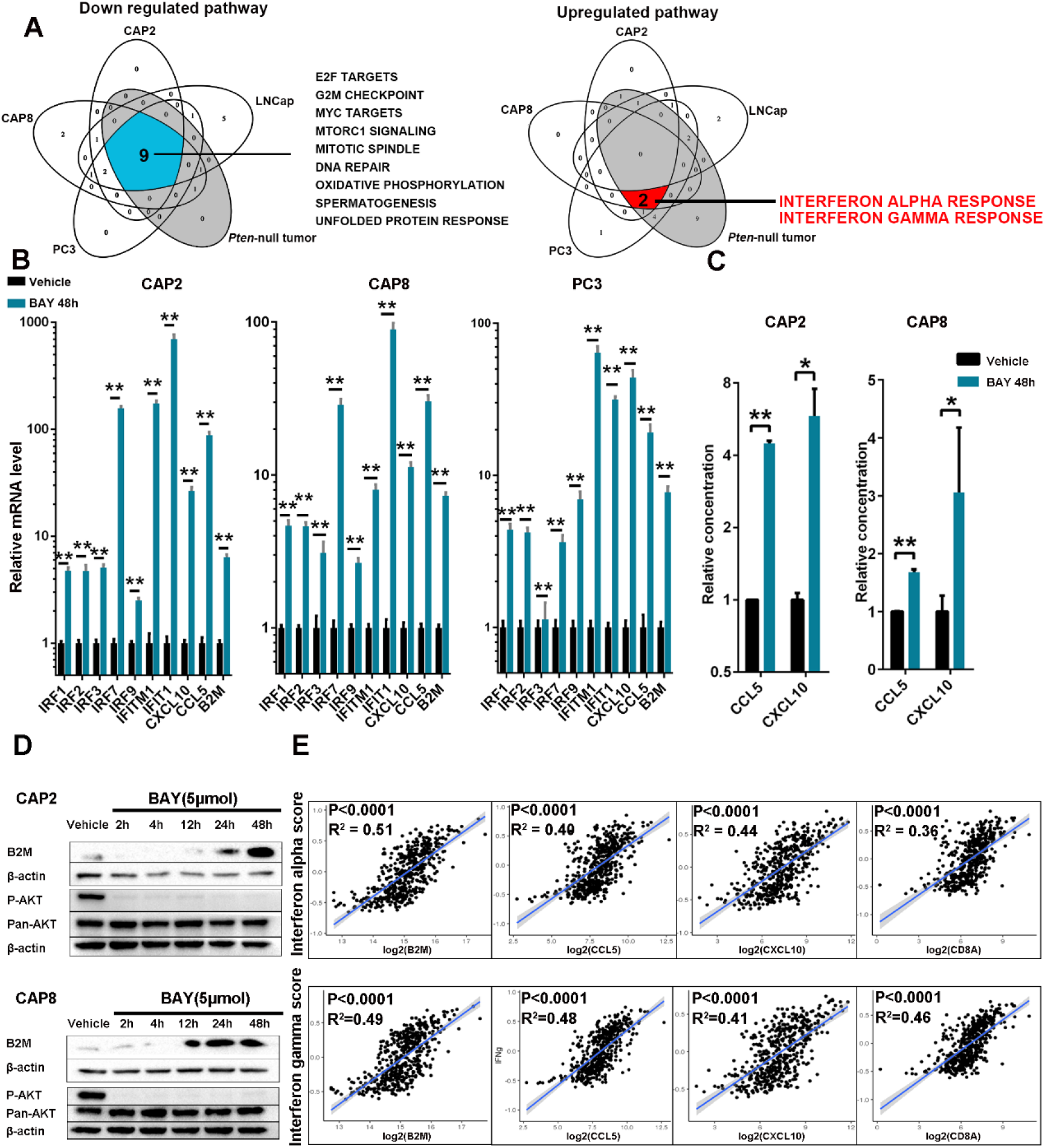
BAY1082439 inhibits the cancer cell-intrinsic immunosuppressive activity. A. PTEN null CAP2, CAP8, PC3 and LNCaP prostate cancer cell lines and *Pten*-null prostate cancer *in vivo* model were treated with vehicle or 5μM BAY1082439 for 48h (for cell lines) or 180mg/kg bullet dose (for in vivo model). RNA-seq and GSEA analyses were performed and commonly down-or up-regulated pathways were presented with p<0.05 as significant. B. RT-PCR analyses showed the relative expression levels of genes related to IFNα/γ pathway, *CCL5/CXCL10* chemokines and *B2M*. C. ELISA measurement for the relative CCL5/CXCL10 secretion levels. PTEN null CAP2 and CAP8 cells were treated with BAY1082439 or vehicle for 48h. Culture supernatant was collected and analyzed as suggested by manufactural recommendation. D. Western blot analyses for P-AKT, Pan-AKT and B2M levels. PTEN null CAP2 and CAP8 cells were treated with BAY1082439 or vehicle for indicated time periods, and cell lysates were analyzed by Western blot using indicated antibodies. E. The positive correlations between IFNα/γ activity scores and CCL5/CXCL10/B2M/CD8A gene expression levels in human prostate cancer tissues. B-D: each experiment was repeated at least 3 times and mean±S.D were presented in B and C with *, p < 0.05, **, p < 0.01.

Quantitative RT-PCR analysis further revealed that IFN-regulated transcription factors, such as *IRF1, IRF2, IRF3, IRF7* and *IRF9,* were upregulated in CAP2/CAP8/PC3 cell lines, but not in LNCaP line, in a time-dependent manner (Fig. 1B; S2A; Supplementary Table 1). Besides the JAK-STAT pathway, the immune modulating effects of BAY1082439 appeared to be PI3K activation-dependent as inducing *PTEN* re-expression in the *PTEN*-null PC3 cells could significantly diminish the effects of BAY1082439 (Fig. S2B).

As a result of IFNα and IFNγ pathway activation, the expressions of *CCL5* and *CXCL10,* chemokines known to have pleiotropic effects on monocytes, NK and T cell migration and activation of T cell proliferations (Harlin et al., 2009; Liu et al., 2011; Lv et al., 2013), were also upregulated (Fig. 1B). ELISA analysis confirmed that BAY1082439 treatment indeed led to increased CCL5 and CXCL10 secretion from CAP2 and CAP8 cells (Fig. 1C). Furthermore, the *β2-microtubulin (B2M)* gene, which is crucial for antigen-presentation to CD8^+^ T cell and causes resistance to anti-PD-1 therapy when deleted (Zaretsky et al., 2016), was up-regulated upon BAY1082439 treatment (Fig. 1B; and Fig. 1D).

The strong positive correlations between IFNα and IFNγ pathway activities and *CCL5*, *CXCL10, B2M* and *CD8A* gene expressions could also be observed in human prostate, lung and melanoma cancers samples (Fig. 1E; S2C-D). Together, these results suggest that BAY1082439 treatment may convert the PTEN-loss induced cancer cell-intrinsic immuno-suppression to immuno-stimulation by upregulating IFNα and IFNγ pathways, increasing cancer cell antigen-presentation, and releasing chemokines to attract immune cells for favorable immune responses.

### Intermittent but not daily BAY1082439 treatment turns *Pten*-null prostate tumors to T cell-inflamed

The potent effect of BAY1082439 in inhibiting cancer cell-intrinsic immunosuppressive activity prompted us to test whether BAY1082439 treatment could turn *Pten*-null prostate cancer to T cell-inflamed and promote T cell-mediated anti-tumor immunity *in vivo*. Unfortunately, daily BAY1082439 treatment (BAY-D), as we performed before (Zou et al., 2018), led to significant decreased tumor- and spleen-associated CD45^+^, CD8^+^ T and CD4^+^ T cell numbers (Fig. 2A-B; Fig.S3A), which could be detrimental for T cell-mediated anti-tumor immunity even if cancer cell-intrinsic immunosuppression could be alleviated.

**Figure 2.**
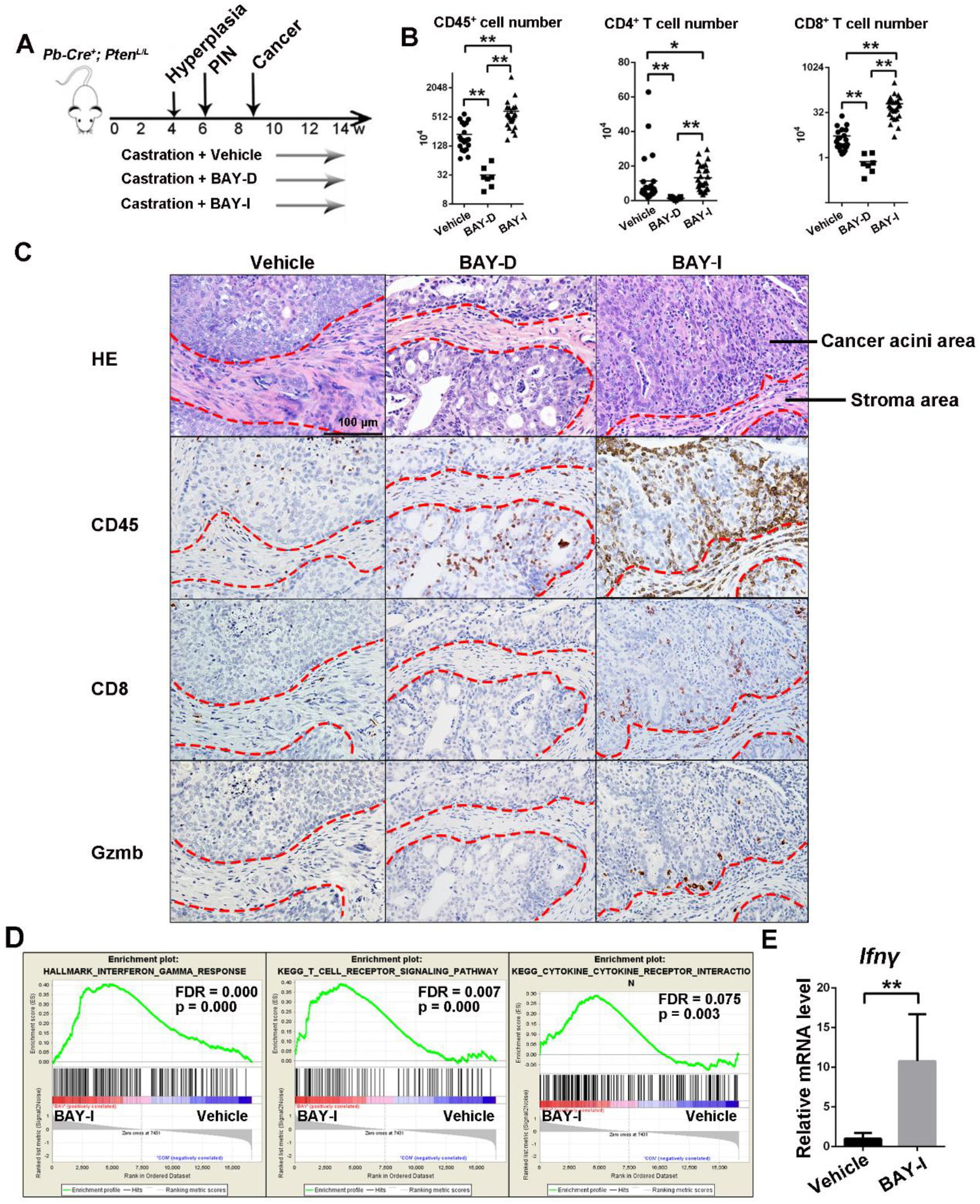
Intermittent but not daily BAY1082439 treatment turns *Pten*-null prostate tumors to T cell-inflamed. A. A schematic illustration of treatment schedules. B. FACS analyses for tumor-associated immune cells. Castrated *Pten*-null mice were treated with vehicle (n=23), BAY-D (n=7) or BAY-I (n=27) for 4 weeks. Tumor tissues were dissociated and weighted, the numbers of tumor-associated CD45^+^, CD4^+^ and CD8^+^ T cells were measured by FACS analysis. Data were presented as whisker blots with medians as the central lines; **, p < 0.01; **, p < 0.01. C. H&E and Immunohistochemistry analyses for immune cell infiltration. Parts of prostate tissues from vehicle or treated cohorts were fixed, and consecutive sections were stained with H&E or antibodies against CD45, CD8 and GZMb. Dashed red lines: the boundaries between cancer acini and stroma areas. D. RNA-seq and GSEA for BAY-I responses. RNAs were extracted from BAY-I treated tumor tissues for RNA-seq analysis. GSEA analysis showed enriched IFN-γ, T cell reporter and cytokine-cytokine receptor interaction signaling pathways in BAY-I treated cohort. E. RT-PCR analysis for increased *Ifnγ* expression in BAY-I treated tumor tissues. Data were presented as mean±S.D; *, p < 0.05, **, p < 0.01.

Since the effects of BAY1082439 on PI3K, tumor growth and IFNα and IFNγ pathways could already be observed 24-48 hours after a bullet dose treatment (Fig. 1A-D; Fig. S2A, S3B), we tested an intermittent (2-day on/5-day off) dosing schedule (BAY-I; Fig. 2A). In contrast to BAY-D, BAY-I treatment led to significantly increased cancer-associated CD45^+^, CD4^+^ and CD8^+^ cell numbers (Fig. 2B-C and data not shown). Importantly, BAY-I, but not BAY-D, treatment could break “the immuno-protective barrier” and promote CD8^+^ and GZMb^+^ cells penetration into cancer acini (Fig. 2C). The effects of BAY-I treatment on T cell activation could also be observed in intact *Pten*-null prostates (Fig. S3C), suggesting that BAY-I could be used in both primary and CRPC settings.

Bulk tumor tissue RNA-seq analysis revealed that BAY-I treatment led to significantly increased IFNγ, T-cell receptor and cytokine-cytokine receptor signaling pathways (Fig. 2D; Supplementary Table 2). RT-PCR analysis further confirmed that the *Ifnγ* expression was increased in BAY-I treatment cohort (Fig. 2E), Finally, the effects of BAY-I treatment on CD8^+^ T cells appeared to be tumor-specific, as CD8^+^ T cell numbers were not changed in other organs, such as the spleen, lung and liver, in the same animals (Fig. S3A and 3D). Thus, intermittent but not daily BAY1082439 treatment could convert the immunosuppressive tumor microenvironment-associated with the *Pten*-null prostate cancer to a T cell-inflamed one.

### Tregs are hypersensitive to BAY1082439 and intermittent BAY1082439 treatment leads to increased tumor-associated CD8^+^/Treg ratios

We next tested the effects of BAY1082439 on different subtypes of T cells by treating freshly isolated CD8^+^, CD4^+^/CD25^+^ helper and CD4^+^/CD25^+^/CD127^low/-^ Treg cells with different concentrations of BAY1082439. As shown in Fig. 3A, Tregs were most sensitive to BAY1082439, followed by T helper, while CD8^+^ T cells were the least sensitive to BAY1082439 (Fig. 3A).

**Figure 3.**
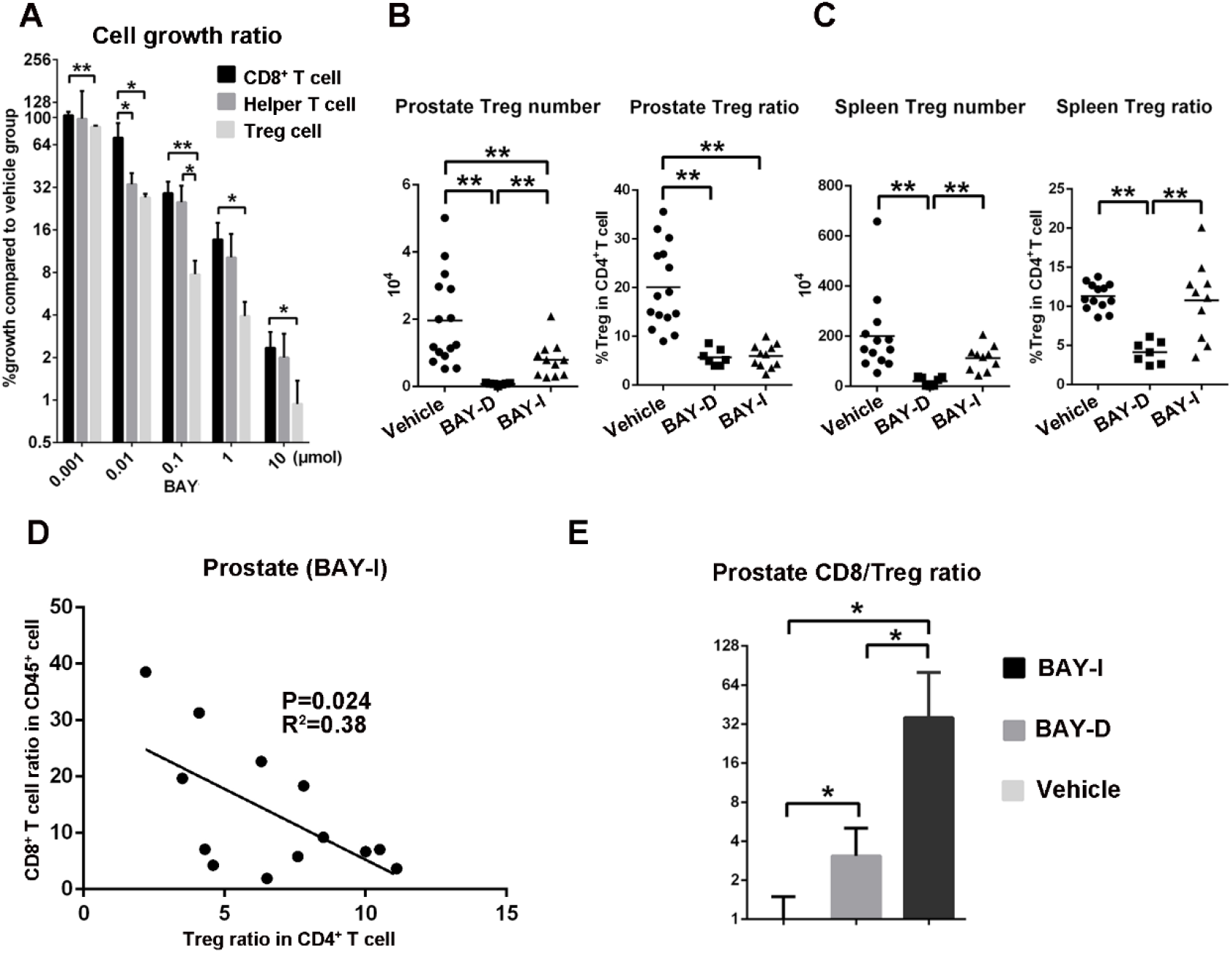
Tregs are hypersensitive to BAY1082439 and intermittent BAY1082439 treatment increases tumor-associated CD8^+^/Treg ratios. A. The differential inhibitory effects of BAY1082439 on T cells. Primary CD8^+^, helper T and Treg cells from spleen of WT mice were FACS sorted, and cultured with different concentration of BAY1082439. This experiment was repeated 3 times and the mean±S.D of the percentages of cell growth as compared to vehicle controls were presented. Data were presented as mean±S.D; *, p < 0.05, **, p < 0.01. B-C. Prostate- and spleen-associated Treg cell numbers and Treg/CD4^+^ T cell ratios in vehicle (n=15), BAY-D (n=7) and BAY-I (n=11) treated cohorts analyzed by FACS. Data were presented as whisker blots with medians as the central lines; **, p < 0.01. D. The negative correlation between CD8^+^/CD45^+^ and Treg/CD4^+^ ratios. E. The CD8^+^/Treg ratios in vehicle, BAY-D and BAY-I treated cohorts analyzed by FACS. Data were presented as mean±S.D.; *, p < 0.05.

We then quantified tumor-associated Tregs in the *Pten*-null model *in vivo*. Both BAY-D and BAY-I treatments led to a significantly decreased total tumor-associated Treg numbers and Treg/CD4^+^ ratios (Fig. 3B). However, BAY-D, but not BAY-I, treatment also led to decreased spleen-associated Treg numbers and Treg/CD4^+^ ratios in the same animals (Fig. 3C), suggesting that intermittent treatment could minimize aberrant immune activation in non-cancerous organs. CD8^+^/CD45^+^ ratios were negatively correlated with Treg/CD4^+^ ratios *in vivo* in the BAY-I treatment cohort (Fig. 3D). Importantly, BAY-I treatment could dramatically increase the intra-tumoral CD8^+^/Treg ratio by 36-fold, as compared to only 3-fold increase in the daily treatment group (Fig. 3E). Therefore, BAY-I treatment could alleviate both cancer cell-associated and Treg-mediated immuno-suppressions, and allow CD8^+^ T cell expansion and activation.

### Intermittent BAY1082439 treatment induces intratumoral CD8^+^ T cell clonal expansion

To study whether the increased CD8^+^ T cells seen in the BAY-I treated prostates were derived from local expansion or recruited from peripheral blood, we treated the *Pten*-null mice with Fingolimod, a S1P inhibitor (S1Pi) that could block lymphocyte egress from hematopoietic organs and lymph nodes (Nikolova et al., 2001). S1Pi treatment effectively depleted T cells in the peripheral blood, but had no effect on BAY-I induced CD8^+^ T cell expansion within the prostates (Fig. 4A), suggesting that increased CD8^+^ T cells after BAY-I treatment were mainly from intratumoral expansion rather than peripheral recruitment. This notion was further supported by BrdU-pulse labeling experiment, as BAY-I treatment doubled the percentage of prostate-associated CD8^+^ cells in cell cycle (Fig. 4B).

**Figure 4.**
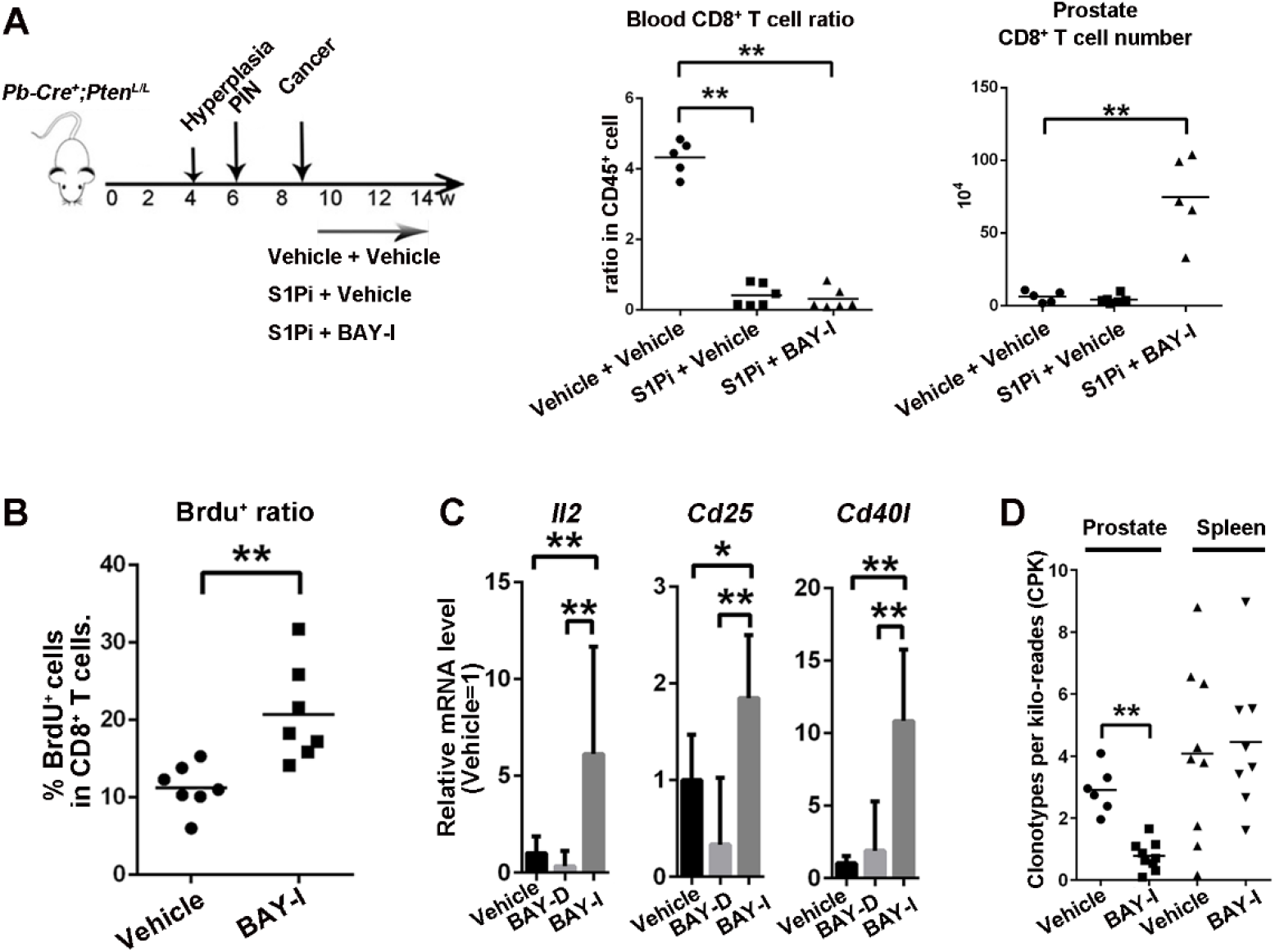
Intermittent BAY1082439 treatment induces tumor-associated CD8^+^ T cell clonal expansion and activation of the IL-2 and TCR signing pathways. A. A schematic illustration of treatment procedures and the ratios of CD8^+^/CD45^+^ in peripheral blood and the prostates. *Pten*-null mice were treated with S1P inhibitor alone (n=5) or in combination with BAY-I (n=6) or vehicle (n=6) for 4 weeks. CD45^+^ and CD8^+^ T cells from peripheral blood or the prostates were sorted and presented as the ratio of CD8^+^/CD45^+^. B. BAY-I treatment induced CD8^+^ T cell proliferation. *Pten*-null mice was treated with BAY-I (n=7) or vehicle (n=7) for 2 weeks, then BrdU labeled (10mg single dose) for 24h before analyzing the percentages of BrdU^+^ cells in CD8^+^ T cell by FACS analysis. C. BAY-I treatment induced CD8^+^ T cell activation. Castrated *Pten*-null mice were treated with vehicle (n=6), BAY-D (n=7) or BAY-I (n=9) for 4 weeks, and tumor-associated CD8^+^ T cells were sorted then analyzed by RNA-seq. The relative expression levels of *Il2, Cd25* and *Cd40l* were presented. D. BAY-I treatment reduced CD8^+^ T cell TCR clonotype diversity. Data were presented as whisker blots with medians as the central lines (for A, B and D) or as mean±S.D (for C); *, p < 0.05, **, p < 0.01.

We next conducted RNA-seq analysis on CD8^+^ T cells isolated from the prostates and found significantly increased the expression of *Il2* as well as T cell activation markers *Cd40l* and *Cd25* expressions in BAY-I treated, but not in BAY-D treated cohort (Fig. 4C; Supplementary Table 2). GSEA pathway analysis revealed enrichment for IL2-STAT5 and T cell receptor signaling pathway in BAY-I but not in BAY-D treated CD8^+^ T cells (Fig. S4). TCR analysis (Li et al., 2016) also revealed a significant decrease in clonotype diversity of tumor-associated, but not in spleen-associated, CD8^+^ T cells upon BAY-I treatment (Fig. 4D). Given BAY1082439 treatment can significantly increase the expression of *B2M* in *PTEN*-null prostate cancer cells (Fig. 1B), this result supports the notion that BAY-I treatment could induce proliferation and activation of tumor antigen-specific T cells.

### Intermittent BAY1082439 treatment-induced anti-tumor immunity is CD8^+^ T cell-dependent

To demonstrate that the BAY-I treatment-induced anti-tumor immunity is CD8^+^ T cell dependent, we crossed the *Pten*-null mice with *Cd8^-/-^* mice (Wai-Ping. et al., 1991) and generated *Pten*-null;*Cd8^-/-^* double knockout mice (DKO). DKO mice developed prostate cancer with similar characteristics as the *Pten*-null mice (data not shown), indicating that CD8^+^ T cells play little role during tumorigenesis in the *Pten*-null mice. Although BAY-I treatment had similar effects on tumor-associated CD4^+^ T cell and Treg cells in the DKO mice (Fig. S5), the increased intratumoral CD8^+^ and GZMb^+^ cells seen in *Pten*-null mice upon BAY-I treatment was nearly completely abolished in DKO mice (Fig. 5A-C). We next investigated BAY1082439 treatment efficacy in the presence or absence of CD8^+^ cells by quantifying cancer cell area in the anterior lobes of the prostates. Comparing to 65% reduction of cancer cell volume in the *Pten*-null prostate, BAY-I treatment of DKO mice showed no significant change (Fig. 5D). Collectively, these results suggest that BAY-I treatment not only directly inhibits *Pten*-null prostate cancer cell growth, but also triggers CD8^+^ T cell-dependent anti-tumor immunity and cancer cell killing effects.

**Figure 5.**
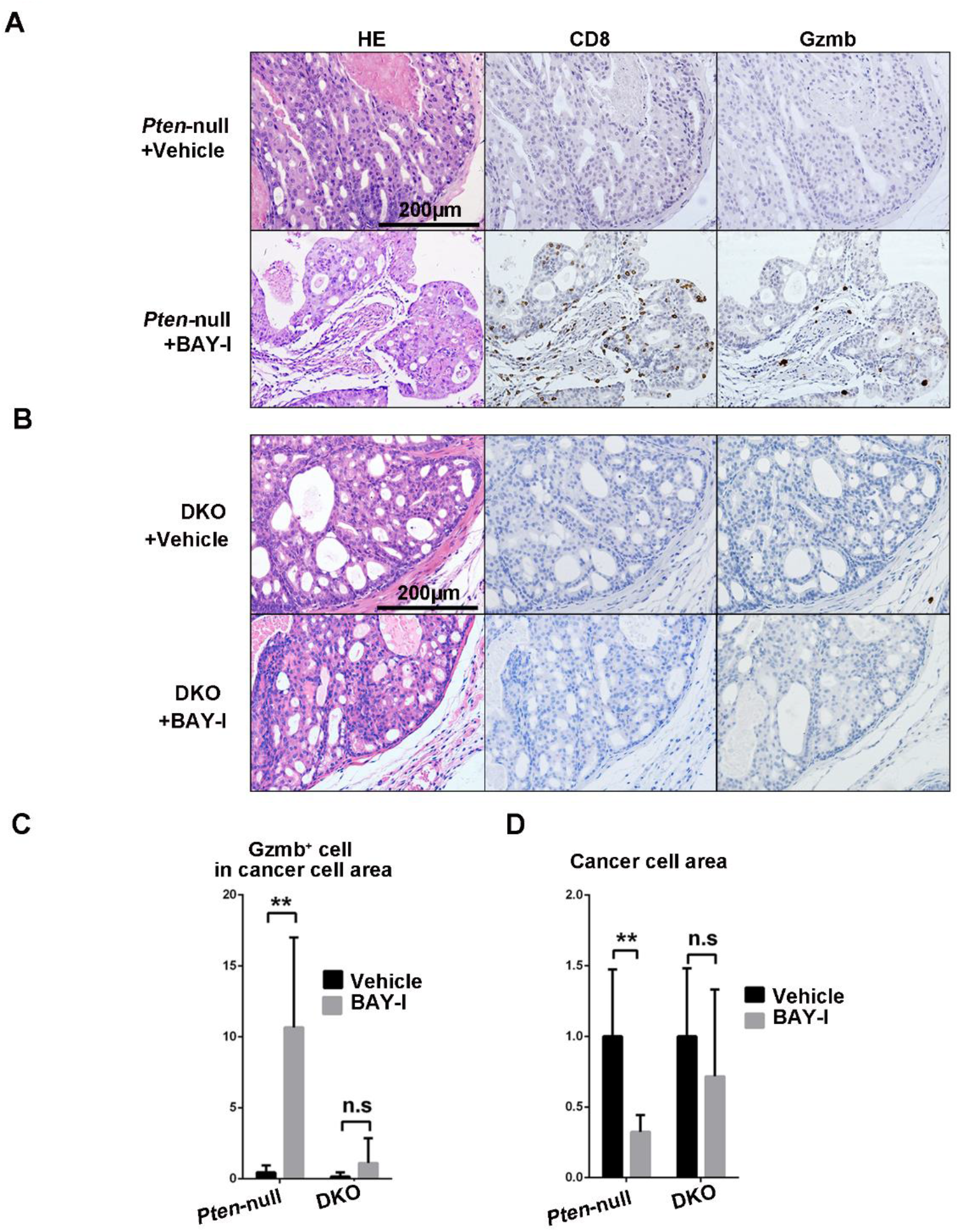
Intermittent BAY1082439 treatment-induced anti-tumor immunity is CD8^+^ T cell-dependent. A-B. BAY-I induced anti-tumor immunity was diminished by genetic deletion of *Cd8. Pten-null* and DKO mice were treated with vehicle (n≥6) or BAY-I (n≥6) for 4 weeks. Tumor tissues were harvested and fixed, and consecutive sections were stained for H&E and antibodies against CD8 and GZMb. C-D. BAY-I treatment efficacies in *Pten*-null or DKO prostates were determined by counting GZMb^+^ cells in cancer areas (C), and calculating cancer cell areas in H&E stained slides (D) for each treatment cohorts. Data were presented as mean±S.D; **, p < 0.01.

### Intermittent BAY1082439 treatment induces prolonged T cell-inflamed phenotype even after drug withdrawal

We next tested whether the BAY1082439 induced T cell-inflamed phenotype persists without continuous drug administration for subsequent combination of ICT. Castrated *Pten*-null mice were treated with vehicle or BAY-I for 4 weeks, then the treatment was stopped for 4 or 10 weeks before the analyses. Interestingly, tumor size and weight were decreased significantly in both 4- and 10-week drug withdrawal groups, as compared to the vehicle or BAY-I group right after the last dose (Fig. 6A). FACS analysis showed that the increased tumor-associated CD8^+^ T cell numbers and CD8^+^/CD45^+^ ratios were well maintained 4- and 10-weeks after drug withdrawal (Fig. 6B), and not affected by S1P inhibitor Fingolimod (Fig. 6C). Importantly, intra-acini CD8^+^ T cell infiltration could be clearly visualized in the prostate tissues 4 and 10-weeks following drug withdrawal (Fig. 6D and data not shown). These results indicate that the BAY-I induced T cell-inflamed phenotype can persist for at least 10 weeks after drug withdrawal, and is likely maintained by intratumoral clonal expansion of tumor specific CD8^+^ T cells, rather than recruitment from the circulation.

**Figure 6.**
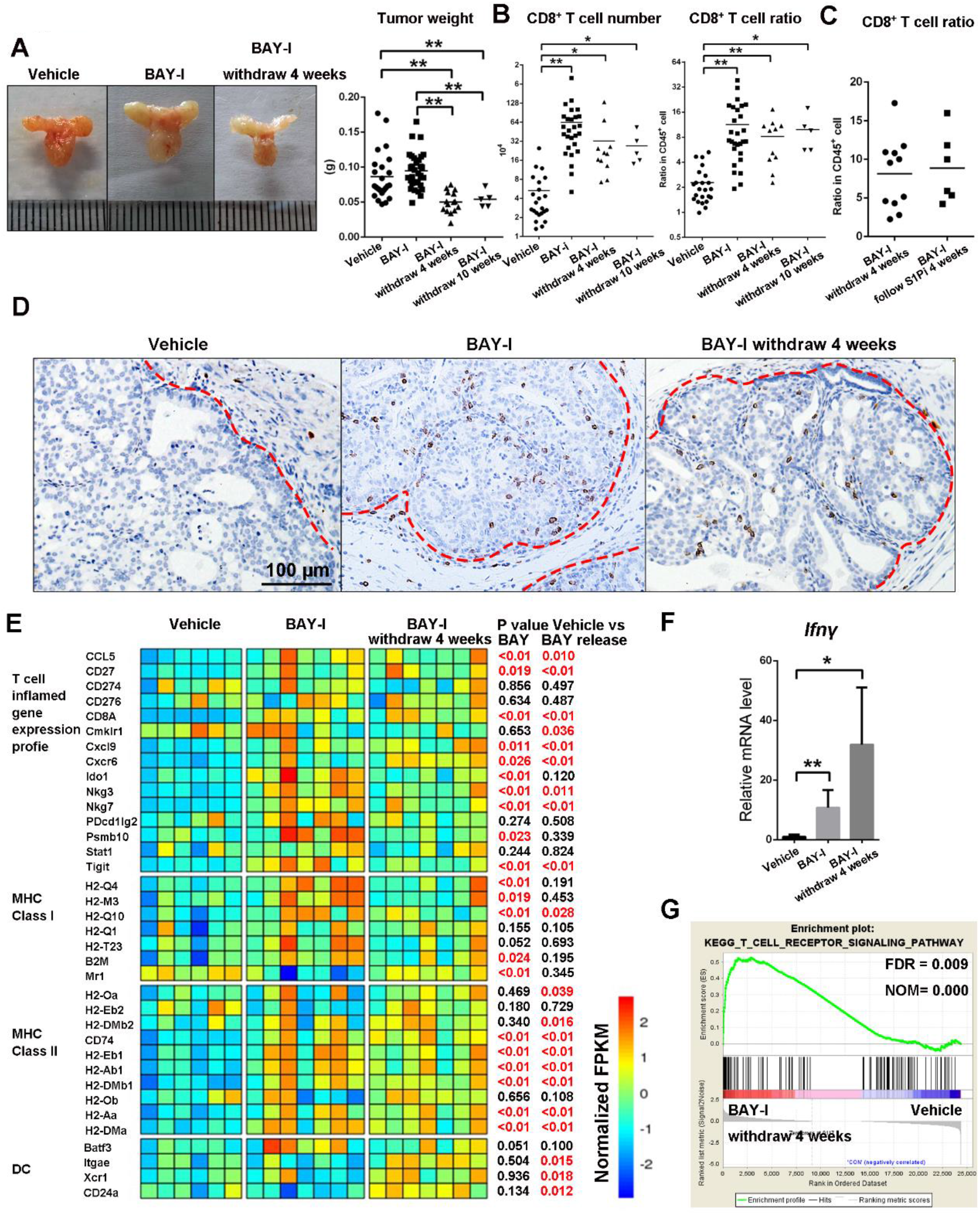
Intermittent BAY1082439 treatment induces prolonged T cell-inflamed phenotype even after drug withdrawal. A-B. BAY-I treatment induced prolonged T cell-inflamed phenotype after drug withdrawal. Castrated *Pten*-null mice were treated with vehicle (n=24), BAY-I (n=32), or BAY-I then drug withdrawal for 4 (n=15) or 10 (n=5) weeks before the analyses. Prostate tumor sizes and weight were presented in (A), and prostate tumor-associated CD8^+^ T cell number and CD8^+^/CD45^+^ ratios were determined by FACS analysis (B). C. BAY-I induced prolonged T cell inflamed phenotype is not influenced by S1Pi. Castrated *Pten*-null mice were treated with BAY-I then drug withdrawal or BAY-I then S1Pi for 4 weeks (n=6). Prostate cancer-associated CD8^+^/CD45^+^ ratios were determined by FACS analysis. D. IHC analysis showed that CD8^+^ T cells remained in the cancer acini 4 weeks after drug withdrawal. Red dash line marks the boundary between cancer acini and stroma areas. E. RNA-seq analysis shows T cell inflamed phenotype after drug withdrawal. RNAs were extracted from tumor tissues in A and B, and RNA-seq analyses were performed. The relative expression levels of indicated genes in each sample were determined and the statistical analysis was performed based on the average of expression levels of each cohort. F. The relative expression levels of the *Ifn-γ* gene were determined by RT-PCR analysis. G. The enrichment of T cell receptor pathway was determined by GSEA based on RNA-seq data in E. A-C, data were presented as whisker blots with medians as the central lines; D, Data were presented as mean±S.D; *, p < 0.05, **, p < 0.01.

RNA-seq analysis of the bulk tumor tissues revealed that BAY-I treatment could increase the T cell-inflamed gene expression profile (Cristescu et al., 2018) and the expressions of MHC class I and II molecules, which were well maintained in 4-week drug withdrawal group (Fig. 6E; Supplementary Table 2). Importantly, the dendritic cell (DC)-associated genes were significantly upregulated in the drug withdrawal group (Fig. 6E). IFNγ gene expression and TCR pathway enrichment were not only maintained but further increased in the drug withdrawal group (Fig. 6F-G). Together, these results demonstrated that BAY-I treatment can prime the tumor and generate a persistent T cell inflammatory environment even in the absence of continued drug administration.

### Intratumoral tertiary lymphoid structures and CD8^+^ memory phenotype-associated with intermittent BAY1082439 treatment

Effective priming of tumor antigen-specific T cells requires secondary lymphoid organs such as lymph nodes (Galluzzi et al., 2018). Since Fingolimod, a S1P inhibitor that could block lymphocyte egress from hematopoietic organs and lymph nodes (Nikolova et al., 2001), did not inhibit intratumoral CD8^+^ T cell activation/expansion induced by BAY-I treatment, we sought alternative mechanisms to explain our findings. In *Pten-*null prostate tumor tissues, we found tertiary lymphoid structures (TLS) with clear B and T cell zones, resembling germinal center morphology (Dieu-Nosjean et al., 2016; Jansen et al., 2019; Sautes-Fridman et al., 2019) (Fig. 7A). When pulse labeled with BrdU, substantial BrdU^+^CD8^+^ double positive cells were found in these intratumoral TLS structures in BAY-I treated group (Fig. 7B), suggesting that these TLS structures may provide the necessary niche for intratumoral CD8^+^ T cell priming and clonal expansion.

These TLS might also account for persistent T cell-inflamed phenotype after drug withdrawal. Comparing RNA-seq data of CD8^+^ T cells revealed increased expression levels of genes associated with T cell activation, effector cytokines, co-inhibitory as well as stimulatory receptors in the BAY-I and drug withdrawal groups but not BAY-D treatment group (Fig. 7C; Supplementary Table 2). The RNA-seq data also revealed a decreased CD62L but increased CD44 and IL7R expression, indicating a shift to memory T cell phenotype. Indeed, FACS analysis demonstrated decreased naïve T cell (CD62L^+^CD44^-^) and increased effector-memory T cell (CD62L^-^CD44^+^) within tumor-associated CD8^+^ T cells in BAY-I treated withdrawal cohort as compared to vehicle cohort (Fig. 7D). Similar to BAY-I group, CD8^+^ T cells isolated from the drug withdrawal group also had decreased clonotype diversity (Fig. 7E).

Cell surface co-expressions of PD-1/CD28 in tumor infiltrating CD8^+^ T cells are positive signs for successful ICT (Daud et al., 2016; Kamphorst et al., 2017; Taube et al., 2014). Thus, we investigated the cell surface expression patterns of PD-1 and other co-stimulatory and inhibitory markers in tumor-associated CD8^+^ T cells in the treatment withdrawal group. FACS analysis revealed that PD-1 and co-stimulatory receptor CD28 and ICOS were upregulated in intratumoral CD8^+^ T cells (Fig. 7F). On the other hand, late exhaustion marker Tim-3 and CTLA-4 were not expressed on the cell surface even through their gene expressions were upregulated (Fig. 7C and F), suggesting that the tumor-associated CD8^+^ T cells were not completely exhausted and may be reprogrammable and activated (Sakuishi et al., 2010). This notion was further backed by the continuously increased expressions of IL2, TNFα/β and IFNγ in tumor-associated CD8^+^ T cell in BAY-I and drug withdrawal groups (Fig. 7C).

We also performed RNA-seq analysis on the EpCAM^+^ cancer cells isolated from BAY-I treatment drug withdawal group. As shown in Fig. 7F, the expression of PD-L1 was significantly increased in BAY-I treated and treatment withdrew cancer cells (Fig. 7G). Taken together, these results indicated that BAY-I treatment can effectively trigger long-term intratumoral clonal expansion and activation of CD8^+^ T cells, most likely via TLS, and reprogram CD8^+^ T cells and cancer cells towards favorable response to ICT.

### Intermittent BAY1082439 treatment paves the way for anti-PD-1 therapy

Upregulated PD-L1 expression in prostate cancer cells upon BAY-I treatment may counteract the cytotoxic activity of CD8^+^ T cells, which provided clear rationale for testing anti-PD-1 combination therapy to achieve maximum tumor killing potential. To test this, we first treated castrated *Pten*-null prostate cancer model with BAY-I for 4 weeks, then dosed with control or PD-1 antibody after drug withdrawal for 4 weeks (Fig. 8A). Anti-PD-1 treatment led to a dramatic cytotoxic effect on otherwise T cell-inflamed tumors, as evidenced by a nearly “hollow” anterior lobe in combinational treatment cohort, which was in sharp contrast to the outcome of BAY-D plus anti-PD-1 group (Fig. 8A, Fig. S6A).

**Figure 7.**
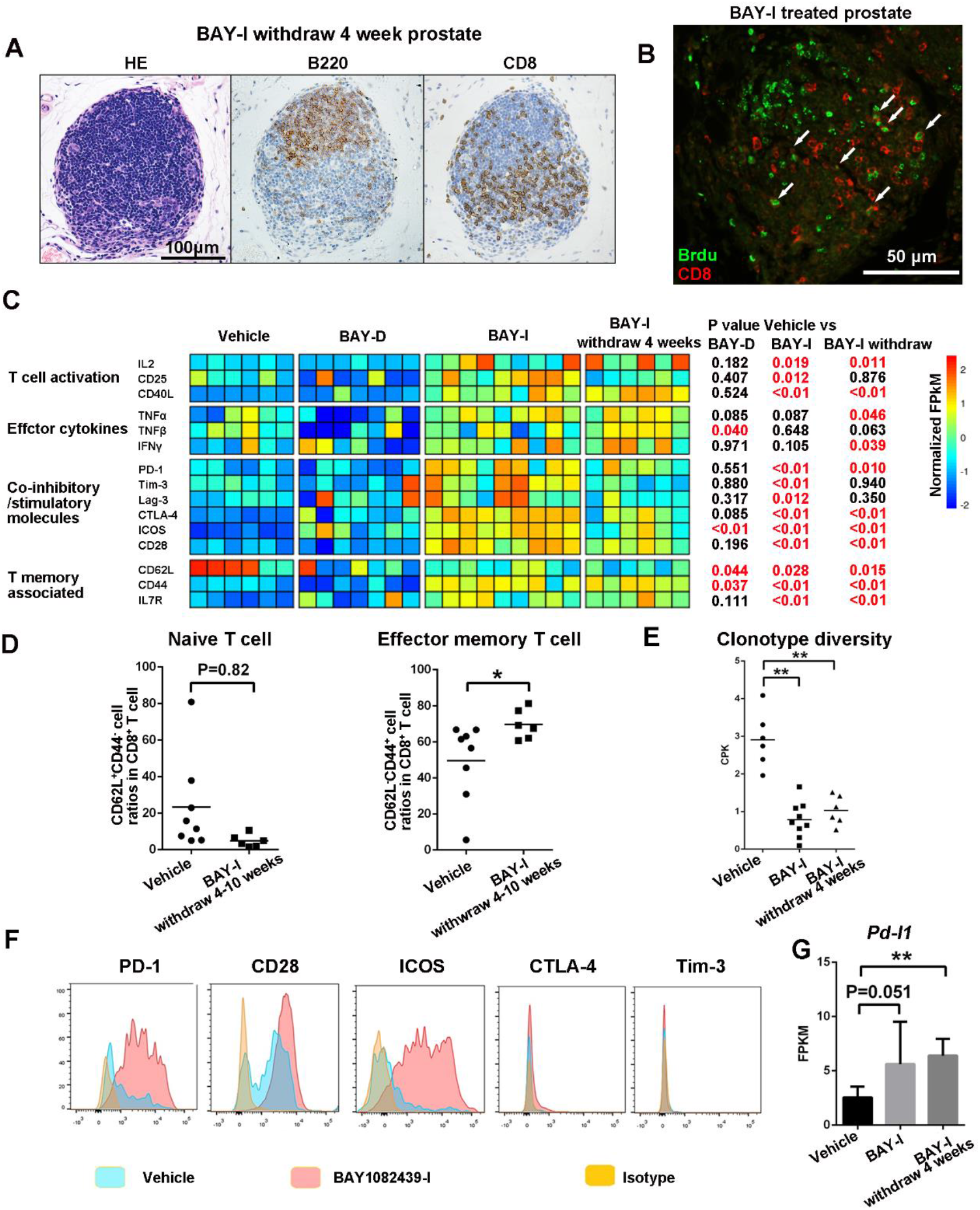
Intratumoral tertiary lymphoid structures and BAY1082439-induced CD8^+^ memory phenotype. A. H&E and IHC analyses showing a representative tertiary lymphoid structure in BAY-I treated prostates. Castrated *Pten*-null mice were treated with 4 cycles of BAY-I then drug was withdrawal for 4-10 weeks. Tumor tissues were fixed, and tertiary lymphoid structures were determined by staining consecutive sections with H&E and anti-CD8 and B220 antibodies. B. CD8^+^ cells within the tertiary lymphoid structures are highly proliferative. Animals in A were pulse-labeled by a bullet dose of BrdU before the analysis. Tumor tissues were fixed and CD8^+^BrdU^+^ double positive cells were visualized by co-Immunofluorescence staining with anti-BrdU and CD8 antibodies. C. BAY-I treated and BAY-I treated then drug withdrew prostate-associated CD8^+^ T cells remained activated. Castrated *Pten*-null mice were treated with vehicle (n=6), BAY-D for 4 weeks (n=7), BAY-I for 4 weeks (n=9) and BAY-I for 4 weeks then drug withdrew for 4 weeks (n=6). Tumor-associated CD8^+^ T cell were sorted by FACS. The relative expression levels of genes associated with T cell activation, effector cytokines, co-inhibitory/stimulatory molecules and T memory were presented based on RNA-seq data. D. Cell surface expression levels of CD44 and CD62L molecules on tumor-associated CD8^+^ T cell were determined by FACS. E. BAY-I treated and BAY-I treated then drug withdrew prostate-associated CD8^+^ T cells remained clonal selected. TCR clonotype diversities were calculated and presented as whisker blots with medians as the central lines; **, p < 0.01. F. Cell surface expression levels of co-inhibitory/stimulatory molecules on tumor-associated CD8^+^ T cell were determined by FACS. G. Increased PD-L1 expressions induced by BAY-I treatment. Lin^-^EPCAM^+^ cancer cells from indicated cohorts were sorted by FACS, and the relative *Pd-l1* expression levels were determined by RNA-seq analysis. Data were presented as mean±S.D; **, P < 0.01.

Although this sequential combination of BAY-I and anti-PD-1 therapies did not significantly alter the numbers of tumor-associated CD8^+^ and Treg cells (Fig. S6B), as compared to BAY-I monotherapy, it did further increased IFNα and IFNγ signaling pathways in tumor tissue (Fig. S6C), GZMb^+^CD8^+^ double positive cell numbers in the remaining tumor areas (Fig. 8E and Fig. S6D), consistent with increased anti-tumor immunity.

**Figure 8.**
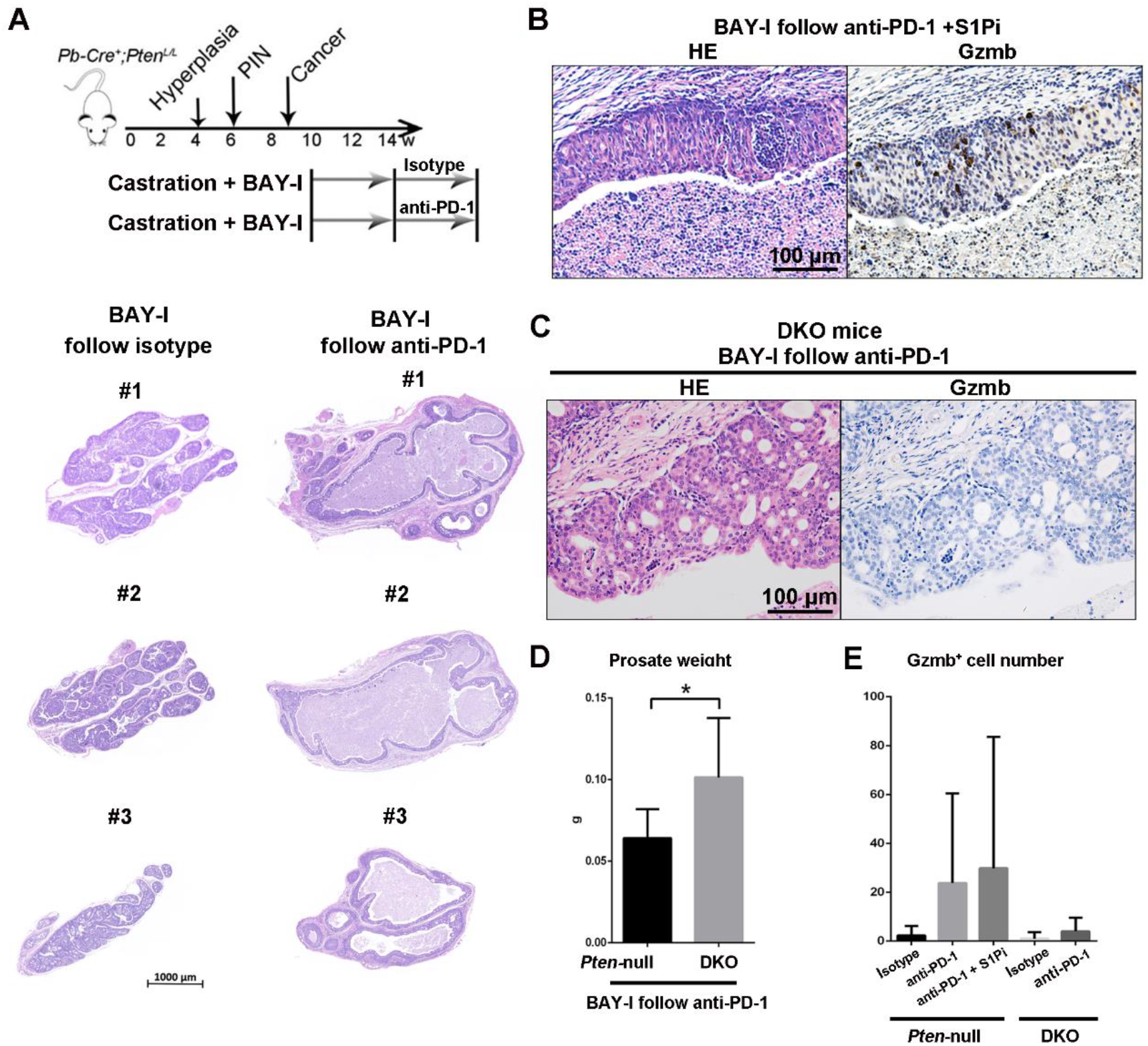
Intermittent BAY1082439 treatment paves the way for subsequent anti-PD-1 therapy. A. A schematic illustration of treatment strategy and low power HE stained images of the anterior prostate lobes. Castrated *Pten*-null mice were treated with BAY-I for 4 weeks followed by drug withdrawal for 4 weeks then isotype antibody (n=5; left) and anti-PD-1 antibody (n=13; right) treatment for 4 weeks. B. The anti-tumor immunity induced by BAY-I and anti-PD-1 combination treatment is not influenced by S1Pi. Similar to A but S1Pi was administrated together with S1Pi for 4 weeks (n=7). H&E and IHC staining for GZMb^+^ cells. C-E. The anti-tumor immunity induced by BAY-I and anti-PD-1 combination treatment is CD8^+^ T cell dependent. The *Pten*-null and *DKO* mice (n≥6 at each group) were treated as indicated in A. Tumor tissues were fixed, weighted, and stained with H&E and anti-GZMb antibody and quantified. BAY-I and anti-PD-1 combination treatment induced anti-tumor immunity was completely diminished by Cd8 deletion (C and E) and the prostate weights from DKO mice were much higher than that of the *Pten*-null prostates (D). Data were presented as mean±S.D; P < 0.05.

The remarkable effects of sequential BAY-I and ICT treatment observed in our study were dependent on intratumoral CD8^+^ T cells, as co-administration of S1Pi with anti-PD-1 antibody did not affect the therapeutic effect of anti-PD-1 antibody (Fig. 8B). To demonstrate the cytotoxic effect of anti-PD-1 antibody treatment depends on CD8^+^ T cells, we treated DKO mice with the same sequential BAY-I and anti-PD-1 combination. The treated DKO prostates had almost no GZMb^+^ cells in the cancer areas (Fig. 8C and E). The prostate weights of DKO mice were also much higher than that of *Pten-* null mice (Fig. 8D). Together, these results demonstrate that a carefully designed dosing schedule for PI3K inhibitor BAY1082439 and sequential administration of targeted and anti-PD-1-mediated ICT can effectively overcome resistance to ICT in a preclinical prostate cancer setting.

## Discussion

Cancer cells have the ability to reshape the surrounding microenvironment for their survival, growth, invasion, and immune escape (Quail and Joyce, 2013; Spranger and Gajewski, 2018). Our study and those by others have demonstrated that PTEN loss or PI3K-AKT pathway activation promote an immunosuppressive microenvironment, resulting in silenced immune response, decreased tumor antigen presentation and decreased CD8^+^ T cell infiltration, which lead to resistance to ICT (Chandrasekaran et al., 2019; Garcia et al., 2014; George et al., 2017; Peng et al., 2016; Sivaram et al., 2019; Wu et al., 2019). We found in this study that PI3K inhibition by BAY1082439 not only inhibits PTEN null prostate cancer cell growth, but also promotes anti-tumor immunity via increased IFNα and IFNγ pathway activities, upregulated expression and secretion of crucial CD8^+^ T cell attracting cytokines and increased antigen presentation (Fig. 1). These results suggest that targeting PI3K signaling in PTEN null cancer cells may be an effective approach to convert immunosuppressive microenvironment into an immunostimulative one to overcome ICT resistance.

Importantly, our study showed that not only the isoform profile but also the dosing schedule of BAY1082439 is critically important for promoting anti-cancer immunity. Intermittent but not daily BAY1082439 treatment can turn *Pten*-null prostate cancers to T cell-inflamed (Figs. 2 and 3). Although both daily and intermittent dosing of BAY1082439 can effectively inhibit PTEN-null prostate tumor cell growth and decrease the number of tumor-associated Tregs, only intermittent dosing schedule can activate the intratumoral cytotoxic CD8^+^ T cells, allow them to undergo clonal expansion, and infiltrate into the cancer acini probably via increased CCL5 or CXCL10 secretion (Figs. 2, 4, 6 and 7). The differential effects of daily vs. intermittent dosing of BAY1082439 on Treg and CD8^+^ T cells are most likely due to the different sensitivities of these immune cells to this anti-PI3Kα/β/δ inhibitor (Fig. 3). As Tregs are most sensitive to BAY1082439, intermittent dosing is sufficient to reduce Treg number and alleviate its immunosuppressive activity, which provides a window for CD8^+^ T cell activation and clonal expansion as indicated by RNA-seq analysis of isolated CD8^+^ T cells (Figs. 3 and 7). Intermittent treatment could also minimize aberrant immune activation in non-cancerous organs, avoiding adverse side-effects (Fig S3). Noticeably, several other target therapy inhibitors have similar inhibitory effects on CD8^+^ T cells, such as androgen receptor antagonists (Pu et al., 2016) and BRAF/MEK inhibitors (Yue et al., 2019). Optimization the dosing schedule of these inhibitors may also improve their therapeutic effects as monotherapies or in combination of ICT.

Intermittent BAY1082439 treatment triggers tumor-specific CD8^+^ T cell activation and expansion, most likely through intra-tumoral TLS, as pre-treating the *Pten*-null CRPC model with S1P inhibitor Fingolimod to block lymphocyte egress from hematopoietic organs and lymph nodes (Nikolova et al., 2001) cannot prevent intra-tumoral CD8^+^ cell activation, clonal expansion and anti-tumor immunity (Figs. 4, 6 and 8). A recent report demonstrates a cloning replacement of tumor-specific T cells following ICT in human basal or squamous cell carcinoma, and suggest that pre-existing tumor-specific T cells may have limited role in ICT (Yost et al., 2019). The different conclusions on the origins of tumor-specific T cells from our study and those by Yost et al may due to the unique characteristics of each cancer type or the specific time windows-associated with each study.

We demonstrate that the CD8^+^ T cells play an essential role in BAY1082439-induced anti-tumor immunity. Importantly, genetically depleting CD8^+^ T cells can completely diminish BAY1082439-induced anti-tumor immunity (Figs. 5 and 8). However, other immune cells may also contribute to the overall therapeutic outcome. We and others have demonstrated the contribution of B cells in CRPC development (Ammirante et al., 2010; Zou et al., 2018) and we have shown that BAY1082439 could significantly inhibit B cells in the *Pten*-null CRPC model (Zou et al., 2018). Similarly, MDSC is increased and DC maturation is decreased in the *Pten*-null model (Calcinotto et al., 2018; Garcia et al., 2014) and the effects of BAY1082439 intermittent treatment on MDSC and DC maturation need further investigation. Together, the superior effects of intermittent BAY1082439 treatment support the idea that drugs that can co-target both cancer cell-intrinsic and microenvironment pathways may have considerably more clinical benefit than single-target drugs.

Recent clinical studies revealed that, depending on the CD8^+^ T cell infiltration levels, solid tumor can be divided into T cell-inflamed “hot tumor” or non-T cell inflamed “cold tumor” (Zou et al., 2016). Treatments that can improve T cell infiltration may augment ICT efficacy (Daud et al., 2016; Jansen et al., 2018; Joyce and Fearon, 2015). Here we show that intermittent BAY1082439 treatment can turn PTEN-null prostate tumor from “cold” to T cell-inflamed (Fig. 2). Intriguingly, once the tumor has become T cell-inflamed, it stays in T cell-inflamed status even after drug withdrawal (Figs. 6 and 7). The CD8^+^ T cells remain in the tumor acini, carry memory T cell signature and are not completely exhausted, while the tumor cells have up-regulated antigen-presentation and high PD-L1 expression (Figs 6–7). This prolonged T cell-inflamed state paves the way for successful anti-PD1 treatment (Fig. 8). Although the detailed mechanisms associated with this prolonged response require further investigation, our study provides a successful pre-clinical case for sequential, instead of simultaneous, anti-PI3K and ICT combination treatment to avoid potential combined toxicity when both drugs are used together.

Our results demonstrate that a carefully designed anti-PI3K treatment, both in its specificity and dosing schedule, to inhibit cancer cell growth while promote anti-tumor immunity, is critically important for successful ICT. Since PTEN loss or PI3K pathway activation is one of the most frequently altered pathways in human cancers, our work may shed light on how to successfully combine anti-PI3K targeted therapy with ICT for broad and long-lasting therapeutic effects on those cancers with PTEN loss or PI3K activation.

## Acknowledgments

The authors thank Dr. Li Wu of Tsinghua university and members of our laboratories for their helpful suggestions and the Flow Cytometry core facility of the National Center for Protein Science at Peking and Tsinghua Universities for the technical assistance. This work was supported by the School of Life Sciences and Peking-Tsinghua Center for Life Sciences at Peking University, as well as awards through the strategic alliance between Peking University and Bayer Pharma to HW.

## Author Contributions

Conception and design: ZQ, NL and HW

Development of methodology: ZQ

Acquisition of data: ZQ, LZ, YZ and WY

Analysis and interpretation of data: ZQ, ZX and HW

Writing, review, and/or revision of the manuscript: ZQ, ZX, NL and HW with help from all authors

Administrative, technical, or material support: CL, NL and HW

Study supervision: HW

## Materials and Methods

### Animals

*Pten*-null mouse model and castration procedure were described previously (Wang et al., 2003). All animal experiment was approved by the Ethics Committee of the Peking University with ID under LSC-WuH-1. *Cd8^atm1Mak^* mice (Wai-Ping. et al., 1991) (CD8^-/-^ mice) was purchased from the Jackson lab (002665) then crossed with the *Pten*-null mice to generate *Pb-Cre^+^;Pten^L/L^;Cd8^-/-^* (DKO) mice.

### Cell culture

The CaP2 and CaP8 cell lines were established and cultured as described previously (Jiao et al., 2007). The PC3 and LNCaP cell lines were purchased from ATCC and the PC3 PTEN inducible cell line are generated as describe (Mulholland et al., 2011) and cultured using Tet-free serum. PTEN re-expression was induced by adding Doxcycline (1ug/ml, sigma D9891) in culture media for 4 days.

### Elisa detection of secreted cytokines

CaP2 and CaP8 cells (1×10^5^) were seed in 6-well plate, cultured in media containing solvent or 5μmol BAY1082439 (dissolved in DMSO with 10mmolTFA as 5mM stock solution). Total cell number was counted using cell counter 48hrs later and supernatant was harvested. Murine CCL5 and CXCL10 concentrations were measured by using Elisa kit R&D MMR00 and Cusabio CSB-E08183h, respectively, according to manufacture recommendations.

### Inhibitors and reagents

BAY1082439 was provided by Bayer AG. S1P inhibitor Fingolimod (FTY720) and Brdu was purchased from MCE.

### Tissue Dissociation, Single-cell Suspension and FACS analysis

Prostates were dissected and photographed, weighted and minced in sterile tissue culture dishes, and subjected to collagenase A (1.5 mg/ml; Roche) and DNase I (0.1 mg/ml; Roche) digestion for 1 h at 37°C with constant agitation. Undigested tissue was removed by passing a 70-μm filter and the total cell number was counted using cell counting chamber. The single-cell suspensions were firstly stained with Fixable Viability Stain 450 (BD) then with fluorescent-labeled antibody against CD45, CD3, CD8, CD4, CD25, PD-1, Tim-3, CTLA-4, CD28, CD44, CD62L or ICOS (Biolegend). FOXP3 was stained by using eBioscience™ FOXP3 Staining Buffer Set. Sorting of each T cell population was performed using BD FACSAria™ III. Different cell population was defined as: CD8^+^ T cells: CD45^+^CD3^+^CD8^+^; CD4^+^ T cells: CD45^+^CD3^+^CD4^+^; Treg: CD45^+^CD3^+^CD4^+^CD25^+^FOXP3^+^. Data was analyzed by using flowjo software. All FACS antibodies used are listed in supplementary table 1.

### In vitro T cell culture

Single-cell suspensions from mice spleen were first stained with Fixable Viability Stain 450 (BD 562274) and CFSE (Thermo C34554) to label live and proliferated cells, followed by fluorescent-labeled antibody against CD45, CD8, CD4, CD25 or CD127 (Biolegend). Viable CD8^+^ T cells (CD45^+^CD8^+^), helper T cells (CD45^+^CD4^+^CD25^-^) and Tregs (CD45^+^CD4^+^CD25^+^CD127^low/-^) from single-cell suspension were sorted by FACS. Cells were seeded at a density of 2.5 × 10^4^/well in 96-well plates pre-coated with 1ug/ml CD3 (Biolegend100339) and 1ug/ml CD28 (Biolegend 102115) antibodies and cultured in 1640 media with 10% FBS, NEAA (Thermo 11140050), 1mM Sodium Pyruvate (Thermo 11360070), 50umol β -ME and 100IU/ml IL2 (Abu-Eid et al., 2014), as well as different concentrations of BAY1082439 or solvent control. Four days later, cells were harvested, live cell numbers were counted by cell counter, and T cell proliferation rate (proliferated cell was determined by CFSE staining) was determined by FACS. Cell growth rate was calculated by the ratio of live, proliferated cell number at each drug concentration vs. solvent control.

### In vivo drug treatment

BAY1082439 was dissolved in 0.1N HCL at 18mg/ml and orally administered. For Single bullet treatment, BAY1082439 was administered at a dose of 180mg/kg/d and mice were analyzed 24h later. For daily treatment, BAY1082439 was administered at a dose of 75mg/kg/d for 4 weeks. For intermittent treatment, BAY1082439 was administered at a dose of 180mg/kg/d in a 2 days on/5 days off manner for 4 weeks. S1P inhibitor (Fingolimod, FTY720) was dissolved in 0.9% NaCl saline solution and orally administered at 1mg/kg/d. Anti-PD-1 (BioXcell, BE0146) or isotype control (BioXcell, BE0089) antibody (200ug per mice) was dosed 3 times per week by i.p. BrdU was dissolved in PBS at 10mg/ml and dosed 100mg/kg by i.p 24h before analysis. BrdU positive cells were analyzed by Flow Cytometry (00-5525) using eBioscience™ BrdU Staining Buffer Set.

### Histology and IHC Analysis

H&E, immunohistochemistry (IHC), and immunofluorescence (IF) staining were performed as described previously (Wang et al., 2003). The antibodies used were listed in Supplementary table 1. To calculate cancer cell area, HE stained slides was scanned, cancer cell area was determined by using CaseViewer software.

### Real time PCR

RNAs were extracted from tumor tissues or cancer cell lines then reverse transcribed using Kit from RNAeasy mini kit (Qiagen 74106) and Vazyme (R223-111 01). Quantitative PCR was achieved using Invitrogen SYBR mix. Primers were listed in Supplementary table 1.

### RNA-seq analysis

For RNA-seq analyses, bulk prostate tissues, sorted LIN^-^EpCAM^+^ prostate cancer cells and cancer-or spleen-associated CD45^+^CD3^+^CD8^+^ T cells were used. RNA was extracted by using RNAeasy mini kit (Qiagen 74106) or micro kit (Qiagen 74004), and cell line or tumor tissue cDNA library was constructed by using NEBNext^®^ Poly(A) mRNA Magnetic Isolation Module (E7490) and NEBNext^®^ Ultra™ RNA Library Prep Kit for Illumina^®^ (E7530). The cDNA library of CD8^+^ T cell and cancer cell was constructed by using SMART-SEQ 2 protocol (Picelli et al., 2014). Sequencing was performed by Illumina – HiSeq PE150.

All RNA-seq data were aligned to the mm10 genome using Tophat (version v2.1.1). Differentially expressed genes were identified by Cuffdiff (version v2.2.1)(Trapnell et al., 2012). FPKM was used for following analysis and comparison. GSEA analysis was performed as software suggested (Mootha et al., 2003; Subramanian et al., 2005). T cell clonotype diversity analysis was performed by TRUST(Li et al., 2016). The pathway activity score was calculated with GSVA(Hänzelmann et al., 2013).

### Analysis of human prostate cancer samples

The relationships between IFNα/γ score and CCL5, CXCL10, B2M and CD8A gene expression in human prostate, lung and melanoma cancer tissues were analyzed by using data from cBioPortal (Cerami et al., 2012; Gao et al., 2013) (Prostate Adenocarcinoma: Firehose Legacy datasets (499 patients); Lung Adenocarcinoma: TCGA PanCancer Atlas datasets (566 patients); Skin Cutaneous melanoma: TCGA PanCancer Atlas datasets (448 patients).

### Statistical Analysis

GraphPad Prism software was used to calculate means and SDs. The Student t test was used to determine statistical significance, and p < 0.05 was considered statistically significant.

## References

Abu-Eid, R., Samara, R. N., Ozbun, L., Abdalla, M. Y., Berzofsky, J. A., Friedman, K. M., Mkrtichyan, M., and Khleif, S. N. (2014). Selective inhibition of regulatory T cells by targeting the PI3K-Akt pathway. Cancer immunology research 2, 1080–1089.

Ammirante, M., Luo, J. L., Grivennikov, S., Nedospasov, S., and Karin, M. (2010). B-cell-derived lymphotoxin promotes castration-resistant prostate cancer. Nature 464, 302–305.

Bilanges, B., Posor, Y., and Vanhaesebroeck, B. (2019). PI3K isoforms in cell signalling and vesicle trafficking. Nat Rev Mol Cell Biol 20, 515–534.

Bonaventura, P., Shekarian, T., Alcazer, V., Valladeau-Guilemond, J., Valsesia-Wittmann, S., Amigorena, S., Caux, C., and Depil, S. (2019). Cold Tumors: A Therapeutic Challenge for Immunotherapy. Front Immunol 10, 168.

Calcinotto, A., Spataro, C., Zagato, E., Di Mitri, D., Gil, V., Crespo, M., De Bernardis, G., Losa, M., Mirenda, M., Pasquini, E., et al. (2018). IL-23 secreted by myeloid cells drives castration-resistant prostate cancer. Nature 559, 363–369.

Carnevalli, L. S., Sinclair, C., Taylor, M. A., Gutierrez, P. M., Langdon, S., Coenen-Stass, A. M. L., Mooney, L., Hughes, A., Jarvis, L., Staniszewska, A., et al. (2018). PI3Kalpha/delta inhibition promotes anti-tumor immunity through direct enhancement of effector CD8(+) T-cell activity. J Immunother Cancer 6, 158.

Cerami, E., Gao, J., Dogrusoz, U., Gross, B. E., Sumer, S. O., Aksoy, B. A., Jacobsen, A., Byrne, C. J., Heuer, M. L., Larsson, E., et al. (2012). The cBio cancer genomics portal: an open platform for exploring multidimensional cancer genomics data. Cancer discovery 2, 401–404.

Chandrasekaran, S., Sasaki, M., Scharer, C. D., Kissick, H. T., Patterson, D. G., Magliocca, K. R., Seykora, J. T., Sapkota, B., Gutman, D. A., Cooper, L. A., et al. (2019). Phosphoinositide 3-Kinase Signaling Can Modulate MHC Class I and II Expression. Mol Cancer Res 17, 2395–2409.

Cheng, J., Huang, Y., Zhang, X., Yu, Y., Wu, S., Jiao, J., Tran, L., Zhang, W., Liu, R., Zhang, L., et al. (2020). TRIM21 and PHLDA3 negatively regulate the crosstalk between the PI3K/AKT pathway and PPP metabolism. Nat Commun 11, 1880.

Cristescu, R., Mogg, R., Ayers, M., Albright, A., Murphy, E., Yearley, J., Sher, X., Liu, X. Q., Lu, H., Nebozhyn, M., et al. (2018). Pan-tumor genomic biomarkers for PD-1 checkpoint blockade-based immunotherapy. Science 362.

Daud, A. I., Loo, K., Pauli, M. L., Sanchez-Rodriguez, R., Sandoval, P. M., Taravati, K., Tsai, K., Nosrati, A., Nardo, L., Alvarado, M. D., et al. (2016). Tumor immune profiling predicts response to anti-PD-1 therapy in human melanoma. J Clin Invest 126, 3447–3452.

De Henau, O., Rausch, M., Winkler, D., Campesato, L. F., Liu, C., Cymerman, D. H., Budhu, S., Ghosh, A., Pink, M., Tchaicha, J., et al. (2016). Overcoming resistance to checkpoint blockade therapy by targeting PI3Kgamma in myeloid cells. Nature 539, 443–447.

Dieu-Nosjean, M. C., Giraldo, N. A., Kaplon, H., Germain, C., Fridman, W. H., and Sautes-Fridman, C. (2016). Tertiary lymphoid structures, drivers of the anti-tumor responses in human cancers. Immunol Rev 271, 260–275.

Dunn., G. P., Sheehan., K. C. F., Old., L. J., and Schreiber., R. D. (2005). IFN Unresponsiveness in LNCaP Cells Due to the Lack of JAK1 Gene Expression. Cancer research.

Galluzzi, L., Chan, T. A., Kroemer, G., Wolchok, J. D., and Lopez-Soto, A. (2018). The hallmarks of successful anticancer immunotherapy. Science translational medicine 10.

Gao, J., Aksoy, B. A., Dogrusoz, U., Dresdner, G., Gross, B., Sumer, S. O., Sun, Y., Jacobsen, A., Sinha, R., Larsson, E., et al. (2013). Integrative analysis of complex cancer genomics and clinical profiles using the cBioPortal. Sci Signal 6, pl1.

Garcia, A. J., Ruscetti, M., Arenzana, T. L., Tran, L. M., Bianci-Frias, D., Sybert, E., Priceman, S. J., Wu, L., Nelson, P. S., Smale, S. T., and Wu, H. (2014). Pten null prostate epithelium promotes localized myeloid-derived suppressor cell expansion and immune suppression during tumor initiation and progression. Mol Cell Biol 34, 2017–2028.

George, S., Miao, D., Demetri, G. D., Adeegbe, D., Rodig, S. J., Shukla, S., Lipschitz, M., Amin-Mansour, A., Raut, C. P., Carter, S. L., et al. (2017). Loss of PTEN Is Associated with Resistance to Anti-PD-1 Checkpoint Blockade Therapy in Metastatic Uterine Leiomyosarcoma. Immunity 46, 197–204.

Hänzelmann, S., Castelo, R., and Guinney, J. (2013). GSVA: gene set variation analysis for microarray and RNA-seq data. BMC bioinformatics 14, 7.

Harlin, H., Meng, Y., Peterson, A. C., Zha, Y., Tretiakova, M., Slingluff, C., McKee, M., and Gajewski, T. F. (2009). Chemokine expression in melanoma metastases associated with CD8+ T-cell recruitment. Cancer research 69, 3077–3085.

Jansen, C. S., Prokhnevska, N., and Kissick, H. T. (2018). The requirement for immune infiltration and organization in the tumor microenvironment for successful immunotherapy in prostate cancer. Urologic oncology.

Jansen, C. S., Prokhnevska, N., Master, V. A., Sanda, M. G., Carlisle, J. W., Bilen, M. A., Cardenas, M., Wilkinson, S., Lake, R., Sowalsky, A. G., et al. (2019). An intratumoral niche maintains and differentiates stem-like CD8 T cells. Nature 576, 465–470.

Jia, S., Liu, Z., Zhang, S., Liu, P., Zhang, L., Lee, S. H., Zhang, J., Signoretti, S., Loda, M., Roberts, T. M., and Zhao, J. J. (2008). Essential roles of PI(3)K-p110beta in cell growth, metabolism and tumorigenesis. Nature 454, 776–779.

Jiao, J., Wang, S., Qiao, R., Vivanco, I., Watson, P. A., Sawyers, C. L., and Wu, H. (2007). Murine cell lines derived from Pten null prostate cancer show the critical role of PTEN in hormone refractory prostate cancer development. Cancer research 67, 6083–6091.

Joyce, J. A., and Fearon, D. T. (2015). T cell exclusion, immune privilege, and the tumor microenvironment. Science 348, 74–80.

Kamphorst, A. O., Wieland, A., Nasti, T., Yang, S., Zhang, R., Barber, D. L., Konieczny, B. T., Daugherty, C. Z., Koenig, L., Yu, K., et al. (2017). Rescue of exhausted CD8 T cells by PD-1-targeted therapies is CD28-dependent. Science 355, 1423–1427.

Kaneda, M. M., Messer, K. S., Ralainirina, N., Li, H., Leem, C. J., Gorjestani, S., Woo, G., Nguyen, A. V., Figueiredo, C. C., Foubert, P., et al. (2016). PI3Kgamma is a molecular switch that controls immune suppression. Nature 539, 437–442.

Li, B., Li, T., Pignon, J. C., Wang, B., Wang, J., Shukla, S. A., Dou, R., Chen, Q., Hodi, F. S., Choueiri, T. K., et al. (2016). Landscape of tumor-infiltrating T cell repertoire of human cancers. Nature genetics 48, 725–732.

Liu, M., Guo, S., and Stiles, J. K. (2011). The emerging role of CXCL10 in cancer (Review). Oncol Lett 2, 583–589.

Lu, X., Horner, J. W., Paul, E., Shang, X., Troncoso, P., Deng, P., Jiang, S., Chang, Q., Spring, D. J., Sharma, P., et al. (2017). Effective combinatorial immunotherapy for castration-resistant prostate cancer. Nature 543, 728–732.

Lv, D., Zhang, Y., Kim, H. J., Zhang, L., and Ma, X. (2013). CCL5 as a potential immunotherapeutic target in triple-negative breast cancer. Cell Mol Immunol 10, 303–310.

Mootha, V. K., Lindgren, C. M., Eriksson, K. F., Subramanian, A., Sihag, S., Lehar, J., Puigserver, P., Carlsson, E., Ridderstrale, M., Laurila, E., et al. (2003). PGC-1alpha-responsive genes involved in oxidative phosphorylation are coordinately downregulated in human diabetes. Nature genetics 34, 267–273.

Mulholland, D. J., Tran, L. M., Li, Y., Cai, H., Morim, A., Wang, S., Plaisier, S., Garraway, I. P., Huang, J., Graeber, T. G., and Wu, H. (2011). Cell autonomous role of PTEN in regulating castration-resistant prostate cancer growth. Cancer cell 19, 792–804.

Nikolova, Z., Hof, A., Baumlin, Y., and Hof, R. P. (2001). Combined FTY720/cyclosporine A treatment promotes graft survival and lowers the peripheral lymphocyte count in DA to lewis heart and skin transplantation models. Transplant immunology 8, 267–277.

Okkenhaug, K., Graupera, M., and Vanhaesebroeck, B. (2016). Targeting PI3K in Cancer: Impact on Tumor Cells, Their Protective Stroma, Angiogenesis, and Immunotherapy. Cancer discovery 6, 1090–1105.

Peng, W., Chen, J. Q., Liu, C., Malu, S., Creasy, C., Tetzlaff, M. T., Xu, C., McKenzie, J. A., Zhang, C., Liang, X., et al. (2016). Loss of PTEN Promotes Resistance to T Cell-Mediated Immunotherapy. Cancer discovery 6, 202–216.

Picelli, S., Faridani, O. R., Bjorklund, A. K., Winberg, G., Sagasser, S., and Sandberg, R. (2014). Full-length RNA-seq from single cells using Smart-seq2. Nature protocols 9, 171–181.

Pu, Y., Xu, M., Liang, Y., Yang, K. T., Guo, Y. J., Yang, X. M., and Fu, Y. X. (2016). Androgen receptor antagonists compromise T cell response against prostate cancer leading to early tumor relapse. Science translational medicine 8.

Quail, D. F., and Joyce, J. A. (2013). Microenvironmental regulation of tumor progression and metastasis. Nature Medicine 19, 1423–1437.

Sakuishi, K., Apetoh, L., Sullivan, J. M., Blazar, B. R., Kuchroo, V. K., and Anderson, A. C. (2010). Targeting Tim-3 and PD-1 pathways to reverse T cell exhaustion and restore anti-tumor immunity. The Journal of experimental medicine 207, 2187–2194.

Sautes-Fridman, C., Petitprez, F., Calderaro, J., and Fridman, W. H. (2019). Tertiary lymphoid structures in the era of cancer immunotherapy. Nature reviews Cancer 19, 307–325.

Sharma, P., and Allison, J. P. (2015). Immune checkpoint targeting in cancer therapy: toward combination strategies with curative potential. Cell 161, 205–214.

Sharma, P., and Allison, J. P. (2020). Dissecting the mechanisms of immune checkpoint therapy. Nat Rev Immunol 20, 75–76.

Sharma, P., Pachynski, R. K., Narayan, V., Flechon, A., Gravis, G., Galsky, M. D., Mahammedi, H., Patnaik, A., Subudhi, S. K., Ciprotti, M., et al. (2020). Nivolumab Plus Ipilimumab for Metastatic Castration-Resistant Prostate Cancer: Preliminary Analysis of Patients in the CheckMate 650 Trial. Cancer cell.

Siegel, R. L., Miller, K. D., and Jemal, A. (2020). Cancer statistics, 2020. CA: a cancer journal for clinicians 70, 7–30.

Sivaram, N., McLaughlin, P. A., Han, H. V., Petrenko, O., Jiang, Y. P., Ballou, L. M., Pham, K., Liu, C., van der Velden, A. W., and Lin, R. Z. (2019). Tumor-intrinsic PIK3CA represses tumor immunogenecity in a model of pancreatic cancer. J Clin Invest 129, 3264–3276.

Spranger, S., Bao, R., and Gajewski, T. F. (2015). Melanoma-intrinsic beta-catenin signalling prevents anti-tumour immunity. Nature 523, 231–235.

Spranger, S., and Gajewski, T. F. (2018). Impact of oncogenic pathways on evasion of antitumour immune responses. Nature Reviews Cancer 18, 139–147.

Subramanian, A., Tamayo, P., Mootha, V. K., Mukherjee, S., Ebert, B. L., Gillette, M. A., Paulovich, A., Pomeroy, S. L., Golub, T. R., Lander, E. S., and Mesirov, J. P. (2005). Gene set enrichment analysis: a knowledge-based approach for interpreting genomewide expression profiles. Proceedings of the National Academy of Sciences of the United States of America 102, 15545–15550.

Taghizadeh, H., Marhold, M., Tomasich, E., Udovica, S., Merchant, A., and Krainer, M. (2019). Immune checkpoint inhibitors in mCRPC - rationales, challenges and perspectives. Oncoimmunology 8, e1644109.

Tang, S., Moore, M. L., Grayson, J. M., and Dubey, P. (2012). Increased CD8+ T-cell function following castration and immunization is countered by parallel expansion of regulatory T cells. Cancer research 72, 1975–1985.

Taube, J. M., Klein, A., Brahmer, J. R., Xu, H., Pan, X., Kim, J. H., Chen, L., Pardoll, D. M., Topalian, S. L., and Anders, R. A. (2014). Association of PD-1, PD-1 ligands, and other features of the tumor immune microenvironment with response to anti-PD-1 therapy. Clinical cancer research: an official journal of the American Association for Cancer Research 20, 5064–5074.

Taylor, B. S., Schultz, N., Hieronymus, H., Gopalan, A., Xiao, Y., Carver, B. S., Arora, V. K., Kaushik, P., Cerami, E., Reva, B., et al. (2010). Integrative genomic profiling of human prostate cancer. Cancer cell 18, 11–22.

Thorpe, L. M., Yuzugullu, H., and Zhao, J. J. (2015). PI3K in cancer: divergent roles of isoforms, modes of activation and therapeutic targeting. Nature reviews Cancer 15, 7–24.

Topalian, S. L., Hodi, F. S., Brahmer, J. R., Gettinger, S. N., Smith, D. C., McDermott, D. F., Powderly, J. D., Carvajal, R. D., Sosman, J. A., Atkins, M. B., et al. (2012). Safety, activity, and immune correlates of anti-PD-1 antibody in cancer. The New England journal of medicine 366, 2443–2454.

Trapnell, C., Roberts, A., Goff, L., Pertea, G., Kim, D., Kelley, D. R., Pimentel, H., Salzberg, S. L., Rinn, J. L., and Pachter, L. (2012). Differential gene and transcript expression analysis of RNA-seq experiments with TopHat and Cufflinks. Nature protocols 7, 562–578.

Tumeh, P. C., Harview, C. L., Yearley, J. H., Shintaku, I. P., Taylor, E. J., Robert, L., Chmielowski, B., Spasic, M., Henry, G., Ciobanu, V., et al. (2014). PD-1 blockade induces responses by inhibiting adaptive immune resistance. Nature 515, 568–571.

Wai-Ping., Fung-Leung., Schilham., M. W., Rahemtulla., l., Kiindig., T. M., Vollenweider., t., Potter., l. J., and Mak’, l. v. E. a. T. W. (1991). CD8 Is Needed for Development of Cytotoxic T Cells but Not Helper T Cells. Cell 65, 443–449,.

Wang, S., Gao, J., Lei, Q., Rozengurt, N., Pritchard, C., Jiao, J., Thomas, G. V., Li, G., Roy-Burman, P., Nelson, P. S., et al. (2003). Prostate-specific deletion of the murine Pten tumor suppressor gene leads to metastatic prostate cancer. Cancer cell 4, 209–221.

Wang, X., Ding, J., and Meng, L. H. (2015). PI3K isoform-selective inhibitors: nextgeneration targeted cancer therapies. Acta Pharmacol Sin 36, 1170–1176.

Watson, P. A., Arora, V. K., and Sawyers, C. L. (2015). Emerging mechanisms of resistance to androgen receptor inhibitors in prostate cancer. Nature reviews Cancer 15, 701–711.

Wu, S., Zhang, Q., Zhang, F., Meng, F., Liu, S., Zhou, R., Wu, Q., Li, X., Shen, L., Huang, J., et al. (2019). HER2 recruits AKT1 to disrupt STING signalling and suppress antiviral defence and antitumour immunity. Nat Cell Biol 21, 1027–1040.

Yost, K. E., Satpathy, A. T., Wells, D. K., Qi, Y., Wang, C., Kageyama, R., McNamara, K. L., Granja, J. M., Sarin, K. Y., Brown, R. A., et al. (2019). Clonal replacement of tumor-specific T cells following PD-1 blockade. Nat Med 25, 1251–1259.

Yue, P., Harper, T., Bacot, S. M., Chowdhury, M., Lee, S., Akue, A., Kukuruga, M. A., Wang, T., and Feldman, G. M. (2019). BRAF and MEK inhibitors differentially affect nivolumab-induced T cell activation by modulating the TCR and AKT signaling pathways. Oncoimmunology 8.

Zaretsky, J. M., Garcia-Diaz, A., Shin, D. S., Escuin-Ordinas, H., Hugo, W., Hu-Lieskovan, S., Torrejon, D. Y., Abril-Rodriguez, G., Sandoval, S., Barthly, L., et al. (2016). Mutations Associated with Acquired Resistance to PD-1 Blockade in Melanoma. The New England journal of medicine 375, 819–829.

Zou, W., Wolchok, J. D., and Chen, L. (2016). PD-L1 (B7-H1) and PD-1 pathway blockade for cancer therapy: Mechanisms, response biomarkers, and combinations. Science translational medicine 8, 328rv324.

Zou, Y., Qi, Z., Guo, W., Zhang, L., Ruscetti, M., Shenoy, T., Liu, N., and Wu, H. (2018). Cotargeting the Cell-Intrinsic and Microenvironment Pathways of Prostate Cancer by PI3Kalpha/beta/delta Inhibitor BAY1082439. Molecular cancer therapeutics 17, 2091–2099.

