## supplemental figures for "Overcoming resistance to immune checkpoint therapy in PTEN-null prostate cancer by sequential intermittent anti-PI3Kα/β/δ and anti-PD-1 treatment"

#### Figure S1

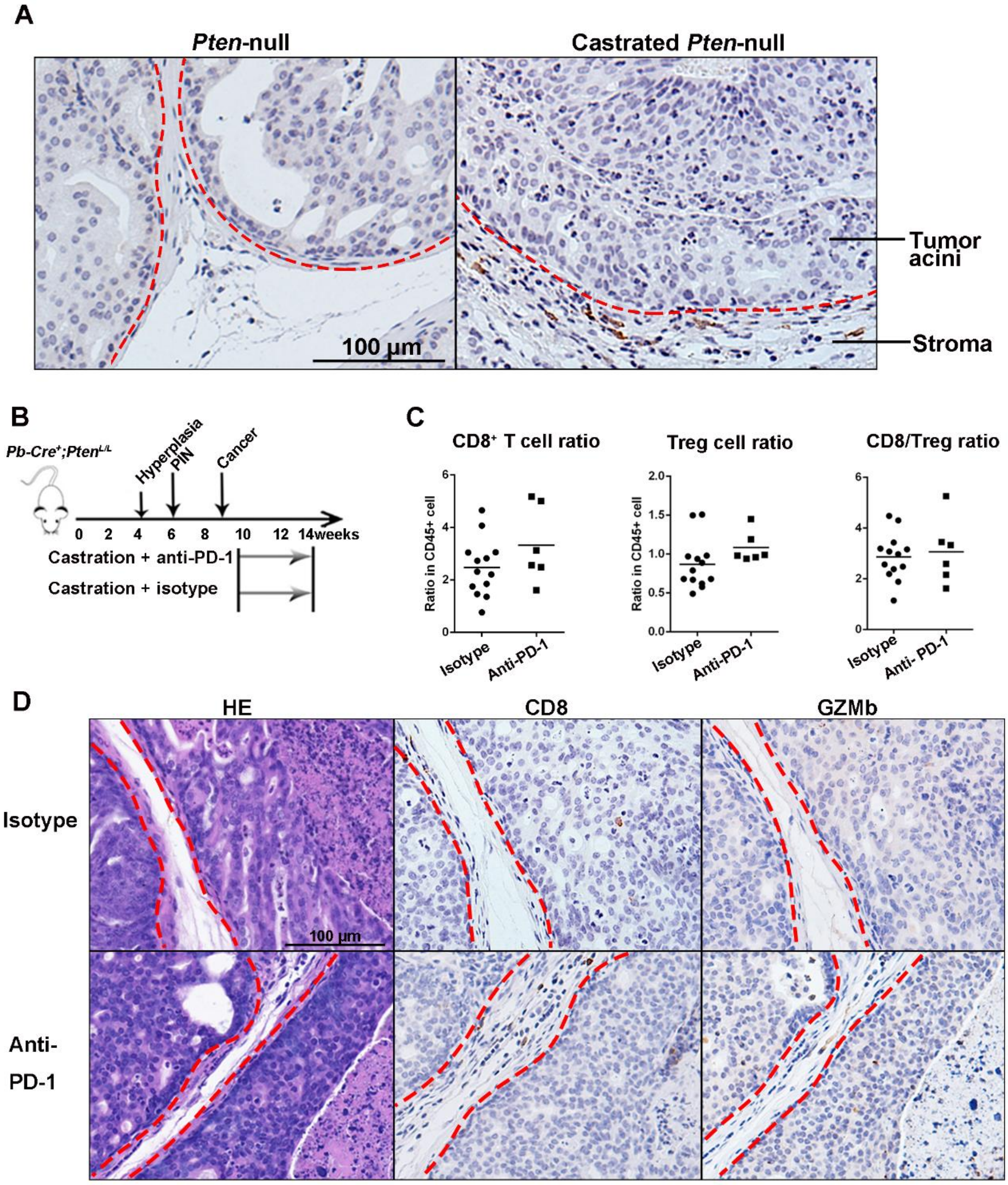

#### Figure S2

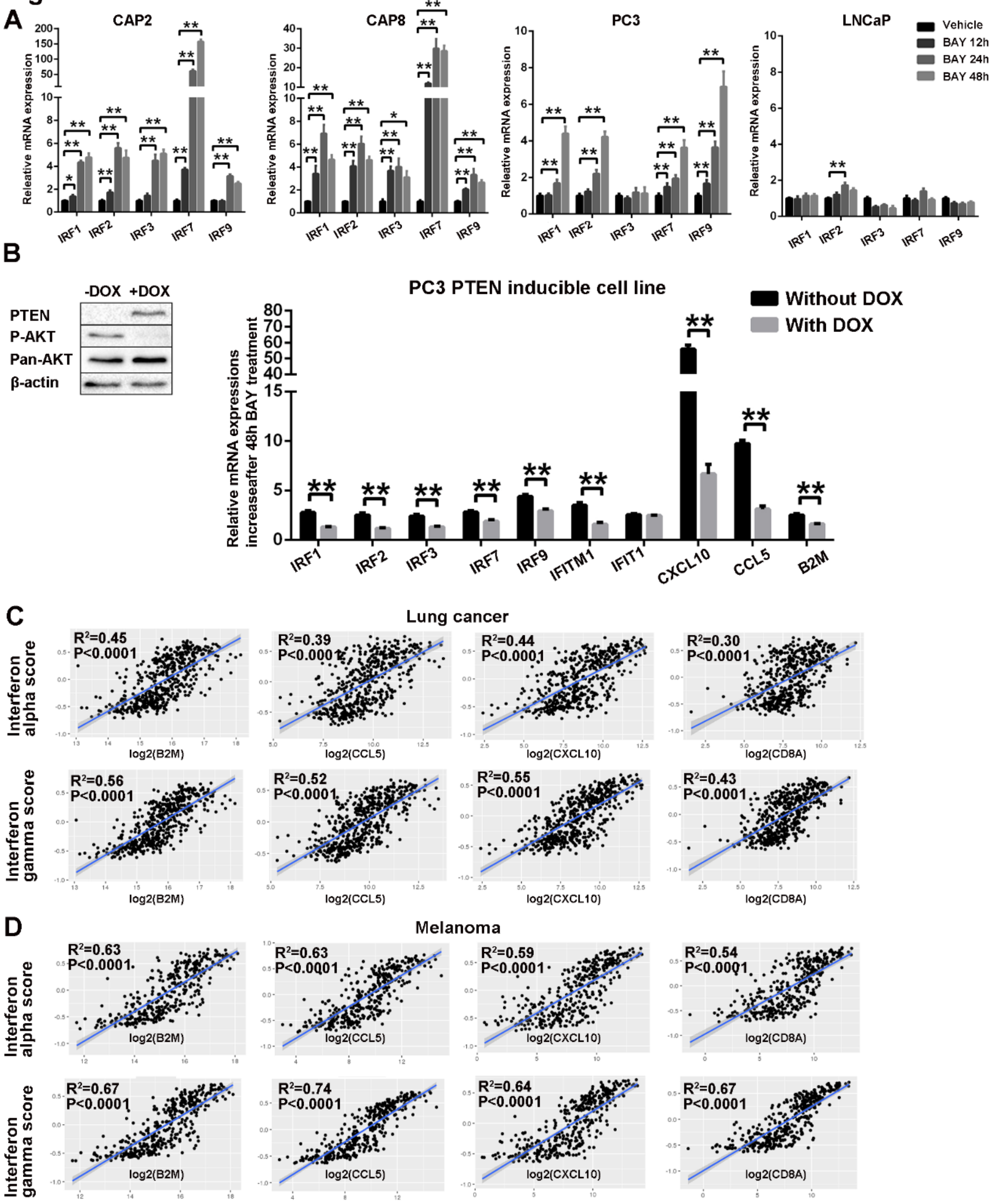

A
CAP2
CAP8
PC3
LNCaP
Vehicle
BAY 12h
BAY 24h
BAY 48h
Releative mRNA expression
IRF1
IRF2
IRF3
IRF7
IRF9
B
-DOX
+DOX
PTEN
P-AKT
Pan-AKT
β-actin
PC3 PTEN inducible cell line
Without DOX
With DOX
Relative mRNA expressions increaseafter 48h BAY treatment
IFITM1
IFIT1
CXCL10
CCL5
B2M
C
Lung cancer
Interferon alpha score
Interferon gamma score
log2(B2M)
log2(CCL5)
log2(CXCL10)
log2(CD8A)
D
Melanoma

### Figure S3

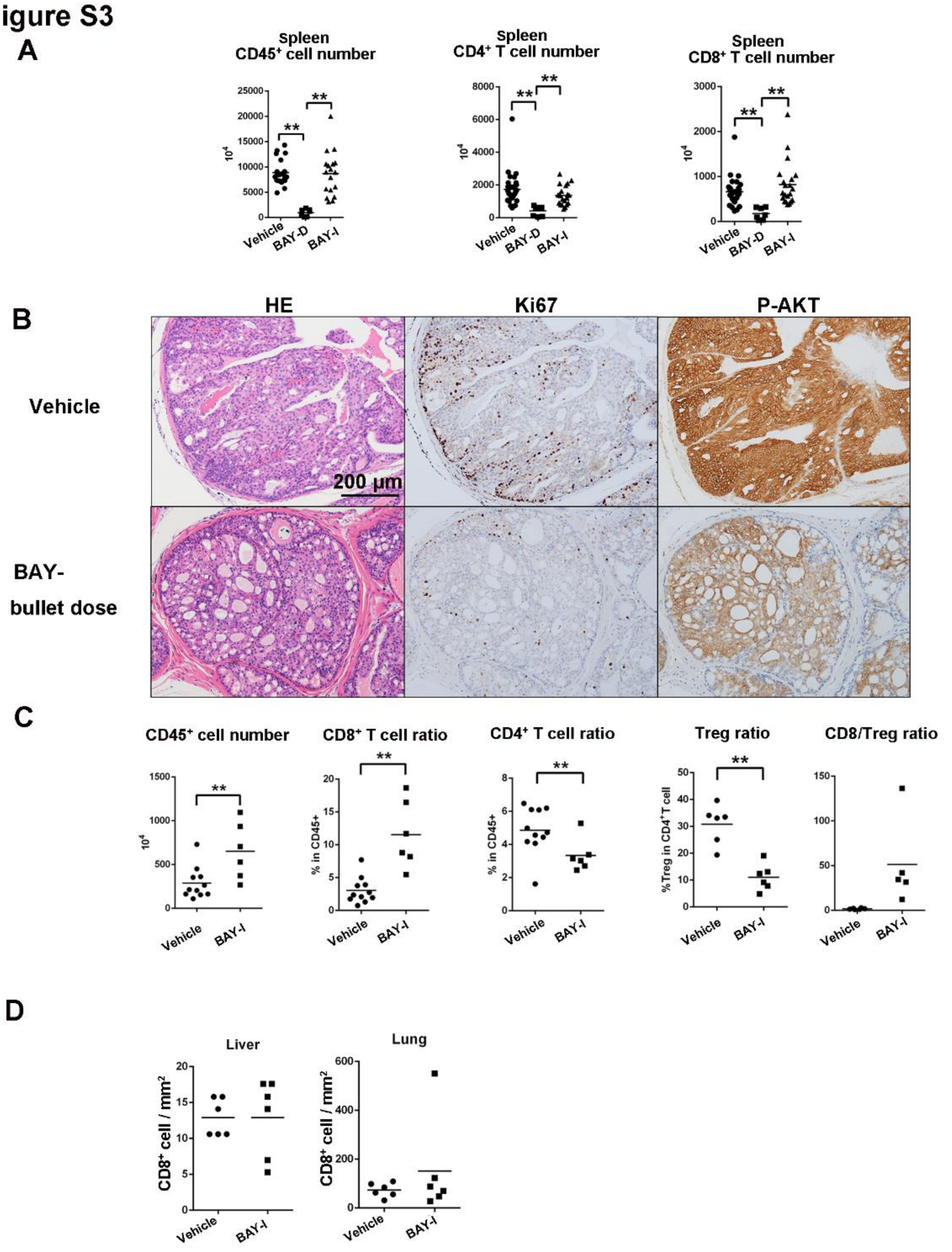

A
Spleen
CD45+ cell number
Spleen
CD4+ T cell number
Spleen
CD8+ T cell number
Vehicle
BAY-D
BAY-I
B
HE
Ki67
P-AKT
Vehicle
BAY-
bullet dose
200 µm
C
CD45+ cell number
CD8+ T cell ratio
CD4+ T cell ratio
Treg ratio
CD8/Treg ratio
% in CD45+
%Treg in CD4+T cell
D
Liver
Lung
CD8+ cell / mm2

### Figure S4

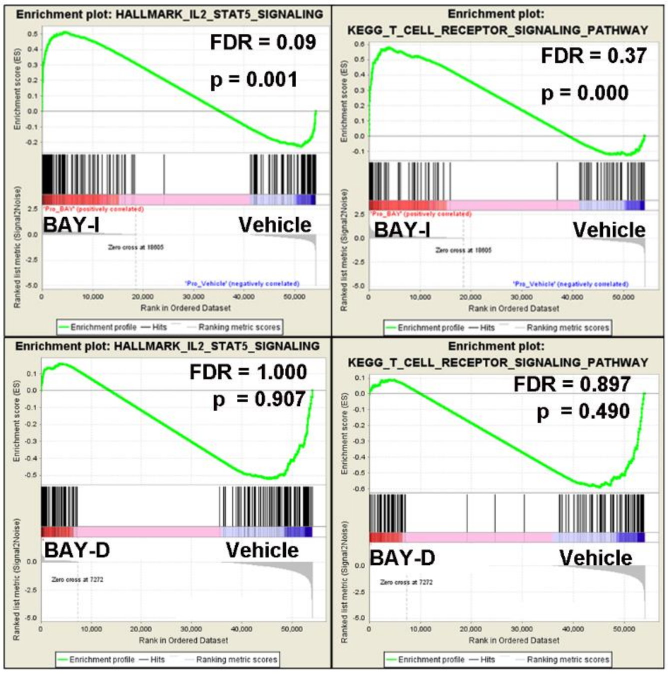

Enrichment plot: HALLMARK_IL2_STAT5_SIGNALING
FDR = 0.09
p = 0.001
BAY-I
Vehicle
Enrichment plot:
KEGG_T_CELL_RECEPTOR_SIGNALING_PATHWAY
FDR = 0.37
p = 0.000
BAY-I
Vehicle
Enrichment plot: HALLMARK_IL2_STAT5_SIGNALING
FDR = 1.000
p = 0.907
BAY-D
Vehicle
Enrichment plot:
KEGG_T_CELL_RECEPTOR_SIGNALING_PATHWAY
FDR = 0.897
p = 0.490
BAY-D
Vehicle

#### Figure S5

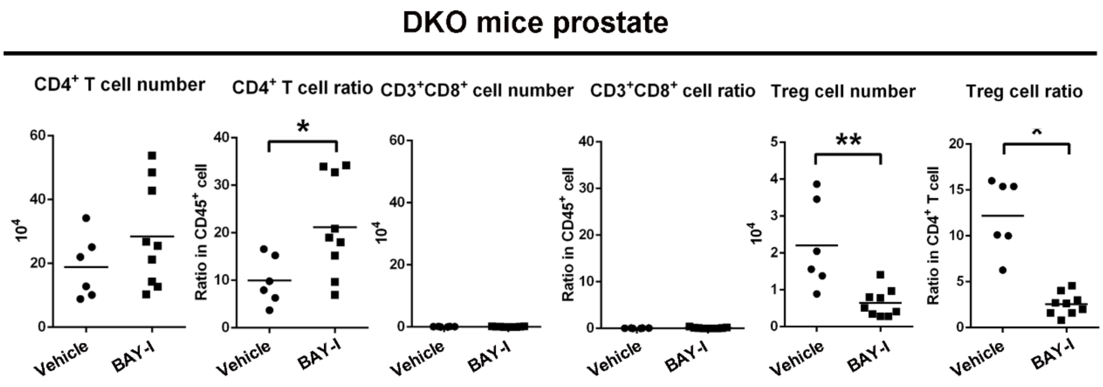

DKO mice prostate
CD4+ T cell number
CD4+ T cell ratio
CD3+CD8+ cell number
CD3+CD8+ cell ratio
Treg cell number
Treg cell ratio
10^4
Ratio in CD45+ cell
Ratio in CD4+ T cell
Vehicle
BAY-I
*
**

Figure S6

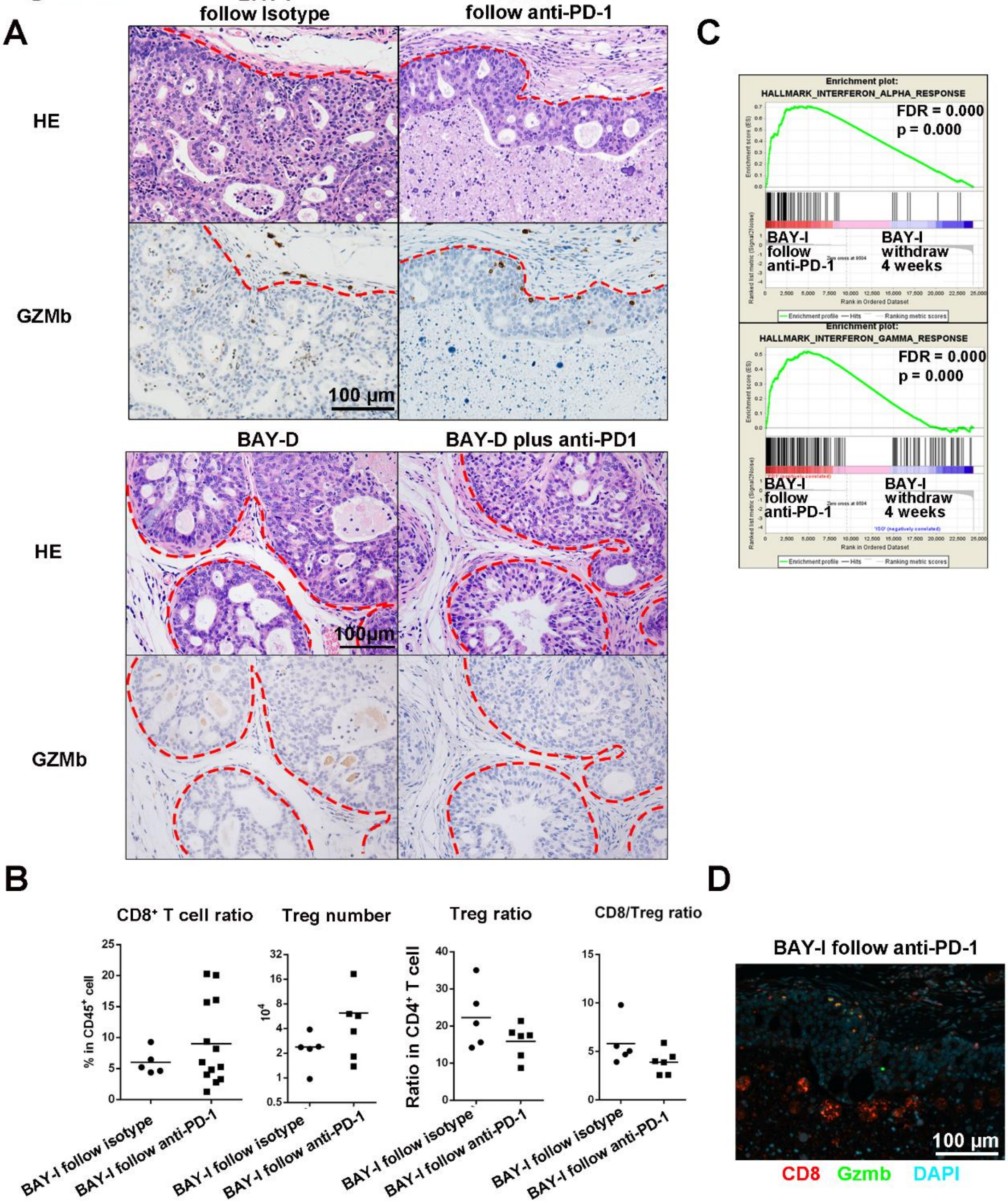
A
BAY-I follow Isotype
BAY-I follow anti-PD-1
HE
GZMb
100 µm
BAY-D
BAY-D plus anti-PD1
HE
100µm
GZMb
B
CD8+ T cell ratio
% in CD45+ cell
Treg number
10^4
Treg ratio
Ratio in CD4+ T cell
CD8/Treg ratio
BAY-I follow isotype
BAY-I follow anti-PD-1
C
Enrichment plot: HALLMARK_INTERFERON_ALPHA_RESPONSE
FDR = 0.000
p = 0.000
BAY-I follow anti-PD-1
BAY-I withdraw 4 weeks
Enrichment plot: HALLMARK_INTERFERON_GAMMA_RESPONSE
FDR = 0.000
p = 0.000
BAY-I follow anti-PD-1
BAY-I withdraw 4 weeks
D
BAY-I follow anti-PD-1
100 µm
CD8 Gzmb DAPI

**Figure S1. *Pten*-null prostate cancers have low T cell infiltration and are resistant to anti-PD-1 monotherapy**

A. Immunohistochemistry (IHC) analysis shows low $CD8^+$ T cell infiltration in primary (left) and CRPC (right) *Pten*-null prostate cancer model.

B-D. A schematic illustration of treatment strategy (B), tumor-associated $CD8^+$/$CD45^+$, Treg/$CD45^+$ and $CD8^+$/Treg ratios after isotype (n=13) or anti-PD-1 (n=6) antibody treatment for 4 weeks (C), and H&E and IHC analyses for infiltrating $CD8^+$ and $GZMb^+$ cells after each treatment (D).

Red dash line marks the boundary between cancer acini and stroma areas. Data in C were presented as whisker blots with medians as central lines.

**Figure S2. BAY1082439 treatment alleviates cancer cell-intrinsic immunosuppressive activity**

A. Time-dependent responses of BAY1082439 (5μM) treatment in four PTEN null prostate cancer cell lines. Relative mRNA expressions were measured by RT-PCR analysis and data were represented as mean±S.D of fold of changes between BAY and vehicle. *, $p<0.05$; **, $p<0.01$.

B. PTEN-dependent BAY1082439 responses in PC3 cells. PTEN was re-expressed in PTEN inducible PC3 cells by doxycycline induction (left). Relative mRNA expressions were measured by RT-PCR analysis without and with PTEN re-expression, and data were represented as mean±S.D of fold of changes between BAY and vehicle. **, $p<0.01$.

C-D. The positive correlations between IFN$\alpha/\gamma$ activity scores and *CCL5/CXCL10/B2M/CD8A* gene expressions in the human lung (C) and melanoma (D) cancer tissues.

**Figure S3. The effects of BAY1082439 treatment on prostate and other organs**

A. The differential effects of BAY-D (n=7) and BAY-I (n=21) treatment on spleen-associated $CD45^+$, $CD4^+$ and $CD8^+$ cells. Data were based on FACS analysis and presented as whisker blots with medians as central lines.

B. *Pten*-null mice were treated with vehicle (n=6) or a bullet dose (n=6) of BAY1082439 (180mg/kg/day for 2 days). 24h later, tumor tissues were fixed and stained for HE, Ki67 or phosphor-AKT (serine 473).

C. BAY-I treatment induced $CD8^+$ T cells expansion and reduced Treg cells in primary prostate cancer. Intact *Pten*-null mice were treated with vehicle (n=11) or BAY-I (n=6) for 4 weeks. $CD45^+$ cell number, $CD8^+/CD45^+$, $CD4^+/CD45^+$, Treg/$CD4^+$ and $CD8^+$/Treg ratios were analyzed by combining FACS with cell counting and data were presented as whisker blots with medians as central lines. **, $P < 0.01$.

D. BAY-I treatment does not influence $CD8^+$ T cells infiltration into other organs. Liver and lung from (A) were fixed and sections were stained with anti-CD8 antibody. $CD8^+$ T cell infiltrations were quantified in vehicle and BAY-I treated cohorts and presented as whisker blots with medians as central lines.

**Figure S4. Differential effect of BAY-D and BAY-I dosing schedules on T cell signaling**

GSEA analysis of RNA-seq data from, Vehicle (n=6), BAY-I (n=9) and BAY-D (n=7) treated $CD8^+$ T cells for IL-2-STAT5 and T cell receptor signaling pathways.

**Figure S5. The roles of $CD8^+$ T cell in BAY-I treatment induced immunity**

*Pten-null;CD8*$^{-/-}$ DKO mice were treated with vehicle (n=6) and BAY-I (n=9) for 4 weeks. $CD4^+$, $CD8^+$ and Treg cell numbers and ratios were analyzed by FACS and cell counting and presented as whisker blots with medians as central lines; *, $p < 0.05$, **, $p < 0.01$.

**Figure S6. Intermittent BAY1082439 treatment paves the way for subsequent anti-PD-1 therapy**

A. Therapeutic effects of BAY-I and BAY-I plus anti-PD-1 (upper panels), and BAY-D and BAY-D plus anti-PD-1 (lower panels) combination treatments evaluated by H&E and IHC analyses. Red dash line marks the boundary between cancer acini and stroma areas.

B. The effect of sequential BAY-I and anti-PD-1 therapies on tumor-associated $CD8^+$ T and Treg cells, as compared to BAY-I monotherapy follow isotype, measured by FACS analysis and presented by whisker blots with medians as central lines.

C. The effect of BAY-I follow anti-PD-1 on tumor tissue IFNα/γ pathway, as compared to BAY-I monotherapy withdraw group.

D. $GZMb^{+}$;$CD8^{+}$ cells detected by co-immunofluorescence analysis in BAY-I follow anti-PD-1.

**Table S1. Primers and key reagents used in this study.**

**Table S2. Gene expression list from RNAseq used in analysis.**
